# Boosting the Predictive Power of Protein Representations with a Corpus of Text Annotations

**DOI:** 10.1101/2024.07.22.604688

**Authors:** Haonan Duan, Marta Skreta, Leonardo Cotta, Ella Miray Rajaonson, Nikita Dhawan, Alán Aspuru-Guzik, Chris J. Maddison

**Affiliations:** Department of Computer Science, University of Toronto, Canada; Vector Institute, Toronto, Canada; Department of Chemistry, University of Toronto, Canada

**Author notes:** Equal Contributions (Haonan Duan, Marta Skreta).

## Abstract

Protein language models are trained to predict amino acid sequences from vast protein databases, while learning to represent proteins as feature vectors. These vector representations have enabled impressive applications, from predicting mutation effects to protein folding. One of the reasons offered for the success of these models is that conserved sequence motifs tend to be important for protein fitness. Yet, the relationship between sequence conservation and fitness can be confounded by the evolutionary and environmental context. Should we therefore look to other data sources that may contain more direct functional information? In this work, we conduct a comprehensive study examining the effects of training protein models to predict nineteen types of text annotations from UniProt. Our results show that finetuning protein models on a subset of these annotations enhances the models’ predictive capabilities on a variety of function prediction tasks. Notably, our model outperforms the search algorithm BLAST, which none of the pre-trained protein models accomplished in our evaluation. Our results suggest that a much wider array of data modalities, such as text annotations, may be tapped to improve protein language models. We host our model checkpoints on https://huggingface.co/h4duan.

## 1 Introduction

We now have a glimpse into the vast expanse of life’s proteome due to advances in sequencing technologies. This treasure trove of data, e.g., the UniProt Knowledgebase [11], has catalyzed the development of large protein language models (PLMs). These models are trained on a pseudo-likelihood objective, and they learn to predict the conditional likelihood of each amino acid residue given its surrounding sequence context [39, 28, 16].

PLMs have demonstrated remarkable versatility, learning protein representations that are useful for a range of modeling and design applications, including remote homology detection [39], prediction of secondary structures [28], enzyme commission number [59] and affinity maturation [19]. Lin et al. [28] offer an intriguing hypothesis to explain this success: the pseudo-likelihood objective encourages PLMs to model the patterns of residue conservation, which requires that PLMs predict the influence of a residue on fitness (sequence co-variation and conservation tends to be predictive of protein fitness [20]). This suggests that PLMs extract functional insights from the patterns encoded in life’s diverse proteome, which mirrors the emergent reasoning capabilities observed in large language models trained on web-scale text databases [53].

However, the pseudo-likelihood objective underpinning PLMs is not without limitations. The relationship between residue conservation in extant sequences and fitness can be confounded by evolutionary and environmental contexts, especially when pooling data from diverse clades [34]. Moreover, raw sequence data fails to capture the wealth of knowledge already derived from direct experimental assays probing protein function. These limitations underscore the potential for developing PLMs that better capture the biological mechanisms governing sequence-function relationships[51]. Where should we look for additional data sources to improve protein language models?

Our key insight is that large sequence databases often contain a wealth of surrounding text that describes structural and functional properties of proteins. While previous works have demonstrated the utility of such annotations in protein representation learning [56, 30, 60, 57], these efforts have been limited in scale and breadth, failing to fully leverage the diverse annotation types available in public databases. For example, You et al. [57] trains DeepText2GO on paper abstracts to predict GO annotations of proteins. The OntoProtein [60] model uses contrastive learning to jointly optimize embeddings of nodes in the GO knowledge graph and protein sequence embeddings. ProtST [56] trains on four types of function annotations with a combination of masked language modelling and contrastive learning losses. Furthermore, there lacks a systematic understanding of how different categories of annotations impact the quality of learned protein representations.

To address these gaps, we compiled the largest public dataset of expertly-curated annotations extracted from the Swiss-Prot subset of UniProtKB [11], encompassing nineteen distinct annotation types. These annotations summarize humanity’s collective knowledge about a variety of key protein properties including domains, family classifications, binding sites, and more. Our results show that fourteen of these annotation types improve PLM representations for function prediction tasks.

To take advantage of this data, we introduce Protein Annotation-Improved Representations (PAIR), a flexible fine-tuning framework that employs a text decoder to guide the fine-tuning process of the encoder. PAIR encourages the learned protein representations to extract information contained within the diverse set of annotations. PAIR’s straightforward and adaptable design allows it to handle any type of text-based annotation, setting it apart from previous, more constrained approaches.

Our results demonstrate that PAIR substantially improves the representational quality of PLMs for function prediction tasks. Using a temporal data split on UniProtKB-Swiss-Prot, we found that PAIR outperforms base PLMs by over 10% across nine diverse function prediction tasks. Notably, PAIR exhibited strong generalization to tasks absent from its training data, indicating an ability to learn generalizable representations across different aspects of protein function.

Perhaps most remarkably, PAIR was the only representation in our experiments that consistently matched or surpassed the performance of a simple-yet-strong BLAST baseline, the *de facto* sequence search algorithm [4]. This performance advantage was particularly pronounced for proteins with low sequence similarity to the training set, suggesting that PAIR captures meaningful information beyond local sequence alignment. Finally, PAIR representations can be pre-computed and stored in vector databases, which allow for more efficient retrieval than a BLAST search [18].

PAIR represents a complementary approach to existing PLM training methods, demonstrating the relatively untapped potential of expert-curated annotations to enhance protein representation learning. We anticipate that our results will inspire future developments in PLMs, encouraging them to look beyond narrowly scoped pseudo-likelihood objectives on sequence datasets.

## 2 Results

### 2.1 PAIR overview

PAIR is a flexible fine-tuning framework to improve the quality of protein representations for function predictions. The core component of PAIR is a transformer-based encoder-decoder model [41], where the encoder takes an amino acid sequence as input and the decoder outputs its associated text annotation. The encoder can be initialized with pretrained protein encoders such as ESM[39] [28] and ProtT5 [16], and the decoder can be initialized with language models trained on biomedical or scientific texts (*e.g.*, SciBert [6]). To model the relationship between the protein sequence and the function annotation, we added a cross-attention module, which makes text tokens attend to amino acids. During training, the parameters of the encoder, the decoder and cross-attention are updated jointly for the model to learn to generate the function annotations from input amino acid sequences. To obtain the final protein representation, we averaged all the amino acid representations from the last layer of the protein encoder.

In order to collect high-quality annotations to train PAIR, we created a database of canonical amino acid sequences and annotations from Swiss-Prot. We downloaded the version of Swiss-Prot published in February 2023. It contains annotations for 569,213 proteins with varying function descriptions, such as name, family and organism. We then parsed 19 types of annotations from Swiss-Prot as shown in Figure 1a. We chose to hold out the annotation type of enzyme commission number (EC) and Gene Ontology (GO) from our training set to evaluate the model’s performance on tasks that it had not been trained on. The details of how we parsed each category can be found in Appendix C. We then generated a train/validation data split based on sequence similarity, as random splitting could bias results if the same sequence (from e.g., different organisms) exists in both training and validation, making it harder to detect overfitting. The details of the split procedure can be found in Appendix B. Our test set was temporally split from the training and validation data, to mimic the real-world setting of annotating newly sequenced proteins using data available at the time. This was done by taking the proteins that had graduated from TrEMBL to the Swiss-Prot subset between February 2023 (2023-02) and January 2024 (2024-01). We visualize this data split in Figure A.

**Figure 1:**
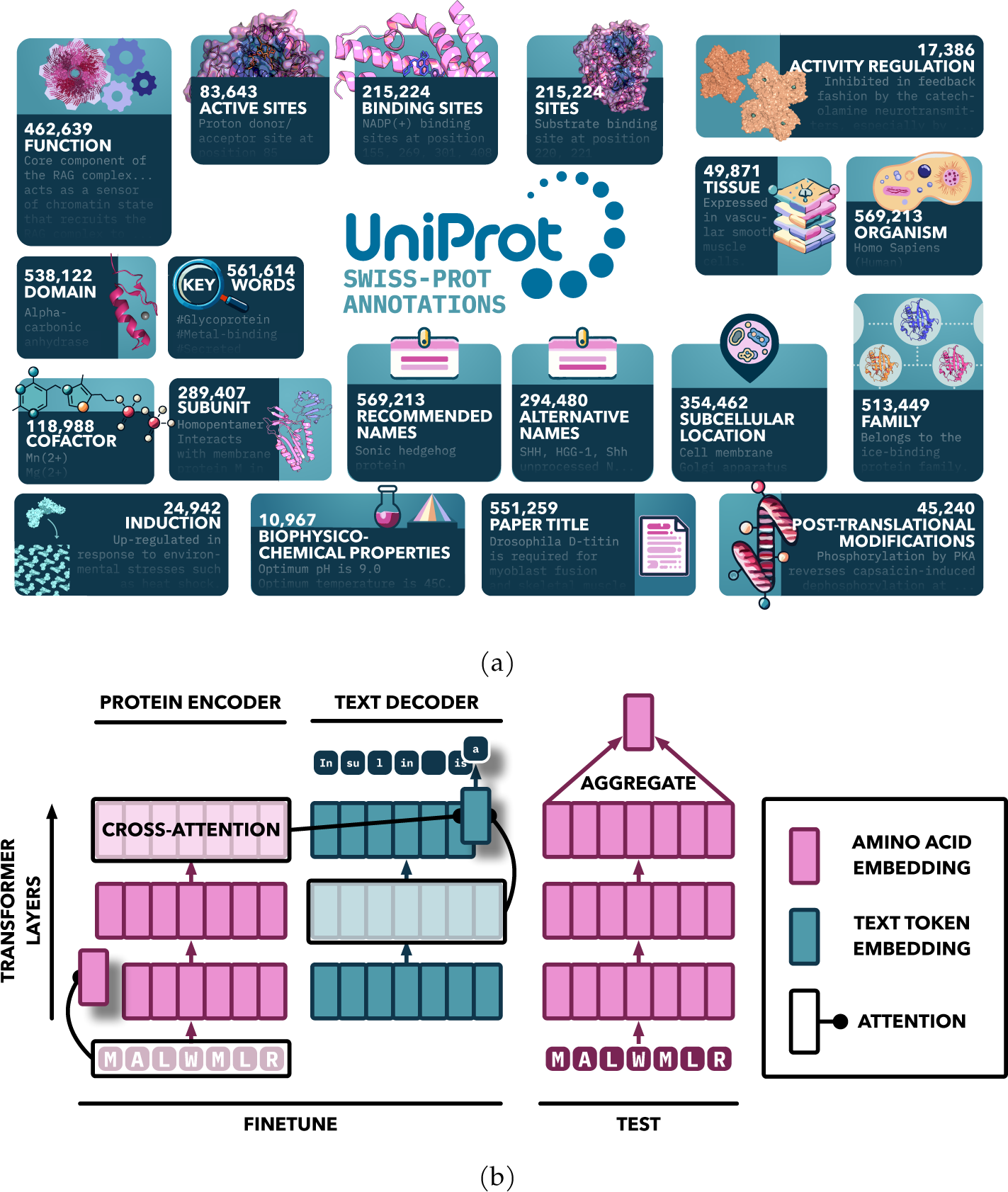
PAIR overview. (a) **Our data.** We parsed 19 types of annotations from Swiss-Prot (February 2023 version). Each box contains the name of the type, the number of proteins having that type of annotations, and an example underneath. (b) **Our model.** PAIR used an attention-based sequence-to-sequence architecture. The encoder and the decoder are initialized by the pretrained models. A randomly-initialized cross-attention is then added between the encoder and the decoder. During training, the encoders, the decoders and the cross-attention are updated jointly to earn to generate the function annotations of the input amino acid sequences. To obtain the final protein representation, we averaged all the amino acid representations from the last layer of the protein encoder.

### 2.2 PAIR’s representation quality relies on the type of annotations

The type of annotation data can strongly influence the quality of the final learned representations. Prior research has not rigorously examined which specific types of annotations are most beneficial for learning high-quality protein representations. To bridge this gap, our initial study systematically evaluated the impact of different annotation data types on the learned representation quality. For a fair comparison, we fine-tuned ESM2-150M on each annotation type separately for one epoch with the same hyperparameters on the training set.

To assess the quality of our representations, we evaluated them on nine function annotation tasks, which we collected from Swiss-Prot. The details of each task can be found in Section 4.2.1. For each task, we performed a nearest neighbour search, a process which is analogous to retrieval methods commonly employed in bioinformatics. Specifically, for a query protein in the validation set, we assigned it the labels of the closest protein(s) in the training set based on Euclidean distance in the embedding space. Nearest neighbour search does not require additional training, which makes it a convenient and fair method for evaluating representation quality. We evaluated on the 10% validation set. To evaluate we used *F*_1_ score for all annotation tasks, except Name (exact match accuracy) and Gene Ontology tasks (weighted *F*_1_).

Figure 2 summarizes the change in performance on different tasks (x-axis) after training on each annotation category (y-axis), compared with the original model ESM2-150M. We also show raw values for all tasks in Table A. Out of the 19 annotation types we collected, training on 14 of them led to improved performance. Among them, *Pfam domain*, *UniProt protein family* and *Recommended name* resulted in the largest performance gains. Moreover, it is remarkable that these performance gains are observed in tasks that the model was not specifically fine-tuned on. This shows PAIR’s ability to learn representations that can generalize beyond the domains that the model was trained on. Finally, this also indicates that, as previously observed in language tasks [33], different protein function tasks require common capabilities.

**Figure 2:**
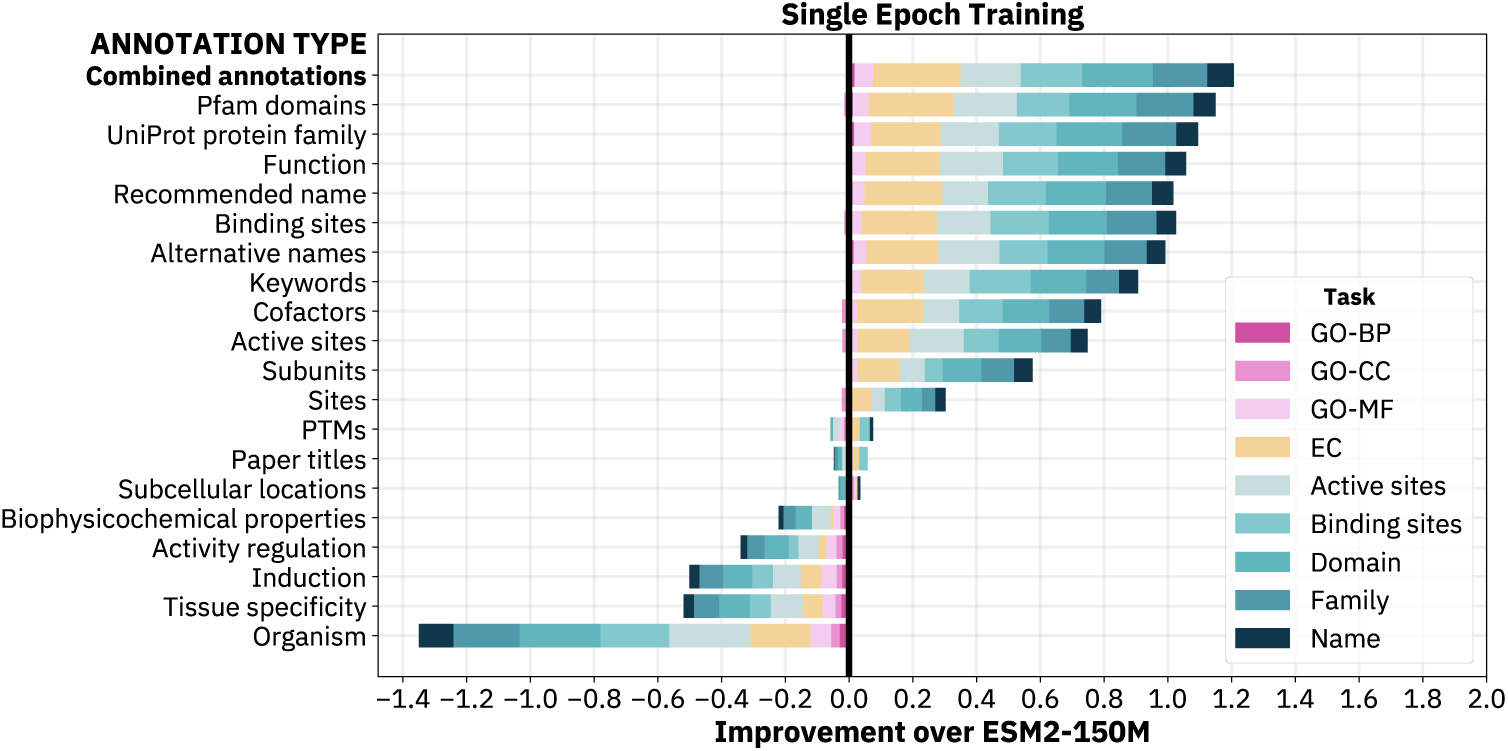
Annotation types play an important role in the quality of learned representations. Each row corresponds to the absolute performance difference on nine tasks between ESM2-150M fine-tuned on an annotation category and baseline ESM2-150M on the validation set. Performance is measured by *F*_1_, weighted *F*_1_, or accuracy depending on the task, which means that the performance difference for each task is in [*−*1, 1]. The black line at 0.0 is the baseline performance of ESM2-150; values greater than 0.0 indicate improved performance for the fine-tuned model, while values less than 0.0 indicate worse performance. We find that training on 14 out of 19 annotations led to improved performance.

On the other hand, we see in Figure 2 that the performance decreased after being fine-tuned on *Organism*, *Tissue specificity*, *Biophysicochemical properties*, *Induction* and *Activity regulation* annotations. We speculate that this performance degradation can be attributed to two reasons. Firstly, *Organism* and *Tissue specificity* represent higher levels of biological organization, and so training to predict them removes information about protein molecular function. Secondly, as shown in Figure 1a, less than 5% of the proteins have annotations for *Biophysicochemical properties*, *Induction* and *Activity regulation*, which may be too little data for the model to learn generalizable representations. Additionally, the *Biophysicochemical properties* of a protein may be too coarse for predicting its function (i.e., knowing that the optimal pH of a protein is 9.0 is not as informative about what the protein does).

Based on the findings above, we removed these five annotation categories from the training set. We then investigated whether it is better to train on the remaining 14 types by combining them all together compared to training on each of them individually. We fine-tuned ESM2-150M on the combined annotations for one epoch and show the performance in the first row of Figure 2. The result indicates that training on all of them yields superior performance compared to training on any single annotation, demonstrating that there is a global synergy between the capabilities required to predict each annotation category.

### 2.3 PAIR improves protein function predictions

We fine-tuned three larger models (ESM2-650M, ESM4541b and ProtT5) with PAIR on the combined 14 annotation types. We denote each fine-tuned model as PAIR(ESM2-650M), PAIR(ESM1b) and PAIR(ProtT5). Training details can be found in Section 4.1.3. Our evaluation is conducted on the temporal test set, which includes all proteins that were added to Swiss-Prot between February 2023 and January 2024 (1631 in total). The results can be found in Table B.

Figure 3a plots the performance difference between each base model and the PAIR-enhanced model on predicting the in-distribution function annotations on the temporal test set. The result demonstrates that PAIR consistently improved upon all those tasks for the three models. Averaged over all model classes, PAIR outperforms the base model by 10.89% for family prediction, 12.53% for name prediction, 10.55% for domain prediction, 8.49% for binding site prediction, 9.84% for active site prediction. ESM2-650M has the largest improvement with 15.10%, 13.56%, 17.31%, 13.55%, 12.41% respectively. Figure 3b evaluates PAIR on the tasks not appearing in its training set: EC and GO. For predicting EC, PAIR improves ESM2-650M, ESM4541b and ProtT5 by 19.39%, 7.47% and 11.89% respectively. PAIR also improves all three subontologies of GO consistently across all models, with average improvements of 7.08%, 12.12%, 5.84% for GO-CC, GO-MF, GO-BP.

**Figure 3:**
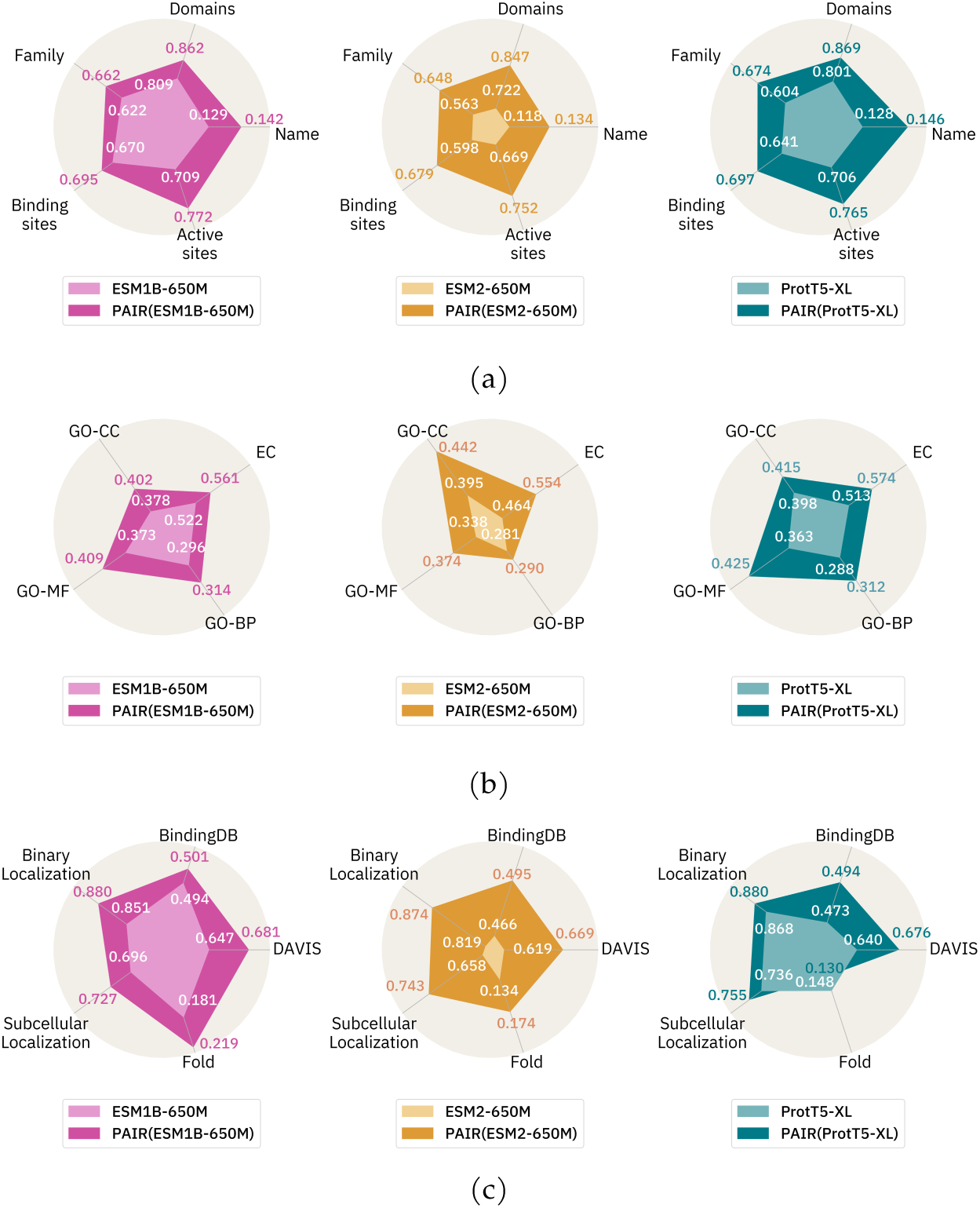
PAIR improves function predictions. PAIR consistently improves upon base models when predicting (a) five function annotation categories in the training set, (b) four function annotation categories *not* in the training set, and (c) five tasks in the external benchmarks. Each spider plot shows the performance of PAIR compared to the base model it was finetuned on. We plot exact performance values for the baseline models in white, PAIR(ESM1b) in pink, PAIR(ESM2-650M) in orange, and PAIR(ProtT5) in blue. To better visualize the performance differences across metrics of different scales, we use Z-score normalization (using the mean and standard deviation across all models) to scale the plots Karamcheti et al. [24]. The scales of the plots are set by the maximum + offset and minimum -offset values within a row, where the offset is set to 0.2 for clearer viewing.

We then evaluated PAIR on five external benchmarks: binary localization [3], subcellular location [3], fold [21], DAVIS [13] and BindingDB [31]. To evaluate the models, we used accuracy for all tasks, except pearson correlation coefficients for DAVIS and BindingDB. A detailed description of these benchmarks can be found in Section 4.2.2. We split the training and test set according to Xu et al. [55], which deliberately lowered the similarity between training and test to evaluate the model’s generalization for remote homologous proteins. For the localization tasks, the training and test set share no more than 30% of sequence similarity. For fold, entire superfamilies are held out in the test set from the training set. Figure 3c shows that PAIR-enhanced model outperforms the original model in all but one set-up.

DAVIS and BindingDB are drug-target interaction (DTI) tasks where the goal is to predict the binding affinity between a molecule and the target protein. We obtained the representations for a molecule-protein pair by concatenating the protein representation with the molecule representation, which we obtained from Text+ChemT5 [10]. We split the dataset so that there is no overlap of protein sequences across training, validation and test. We repeated this random split 50 times and report the average. Figure 3c shows that PAIR improves the DTI predictions for all models over the two datasets.

### 2.4 PAIR outperforms BLAST

We compared the retrieval performance of all PLM representations (with and without PAIR finetuning) to the *de facto* algorithm for retrieval based on sequence similarity in bioinformatics, BLAST (basic local alignment search tool [4]). First, we find that the base models (ESM and ProtT5) perform significantly worse than BLAST, as shown in Figure 4a. Even when compared with the best-performing base model ESM4541b, BLAST still outperforms it on eight of the nine tasks. In the case of EC predictions, BLAST surpassed ESM4541b by a substantial margin of 10.54%. However, after fine-tuning with our approach, PAIR(ProtT5) emerges as the sole model that either equals or surpasses BLAST’s performance across the evaluated tasks. As illustrated in Figure 4a, PAIR(ProtT5) outperformed BLAST on six tasks and achieved comparable results on the remaining three. In addition, we compared PAIR with ProtST [30]. ProtST also provides a framework to fine-tune the protein encoders on text annotations. Figure 4a shows that PAIR consistently outperforms ProtST across tasks given the same pretrained model.

**Figure 4:**
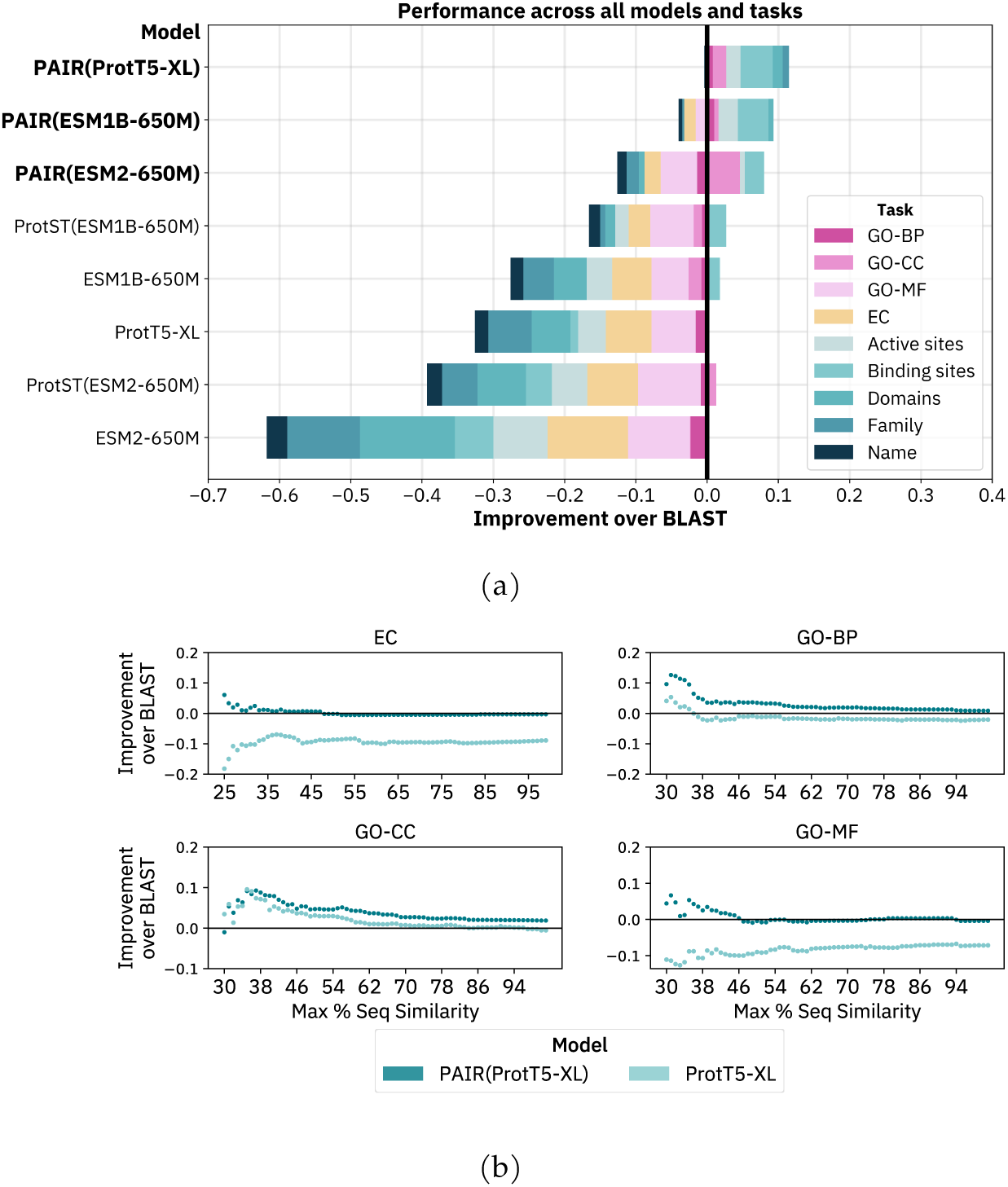
Comparison with BLAST. (a) **PAIR is on par with or superior to BLAST in annotating uncharacterized proteins.** Each row represents the performance of individual models across nine functional annotation tasks on the test set, benchmarked against BLAST. Bars extending to the right imply an improvement over BLAST for the corresponding task, whereas bars extending to the left indicate inferior performance compared to BLAST. Performance is measured by *F*_1_, weighted *F*_1_, or accuracy. The results show that BLAST outperforms all base models on almost all downstream tasks. However, after fine-tuning with our approach, PAIR(ProtT5) emerges as the sole model that either equals or surpasses BLAST’s performance across all the evaluated tasks. (b) **PAIR demonstrates larger advantages over BLAST when predicting the proteins with low sequence similarity.** The x-axis represents the maximum sequence similarity between a test sample and the training set. The y-axis depicts the performance difference between the model and BLAST, where positive values indicate superior performance compared to BLAST. The lighter blue line corresponds to the original ProtT5 model, while the darker blue line represents PAIR(ProtT5). The results demonstrate that our approach, PAIR, exhibits superior performance when predicting the test samples with low sequence similarity to the training data.

We examined how performance of PAIR and BLAST varied with test sequence similarity to the training set. Our results in Figure 4b indicate that PAIR exhibits greater gains in performance when the sequence similarity to the training set is low. This finding is particularly promising because low similarity samples pose a greater challenge for sequence search methods, and this fact suggests that PAIR learns information complementary to BLAST —and possibly other pairwise alignment methods.

We evaluated our approach against BLAST on the three protein prediction external benchmarks. Our method demonstrated substantially superior performance compared to BLAST (binary localization: 73.76, subcellular location: 53.73, fold: 2.92). We attribute the poor performance of BLAST to the relatively small size of the other datasets and minimal overlap between their training and test sets, which is a scenario where BLAST faces challenges. This suggests that PAIR generalizes better to previously unseen data when training examples are limited.

Retrieval over vector databases can be more computationally efficient than BLAST [18]. Because PAIR representations can be pre-computed and stored in a vector database, PAIR opens the door to a highly efficient and performant retrieval algorithm for protein prediction tasks.

### 2.5 PAIR can better capture enzyme function similarity

Understanding enzyme function is important for applications such as pharmaceutical development and chemical synthesis. However, computational prediction of enzyme function is challenging, especially for unstudied or poorly understood enzymes. Traditional computational methods struggle with enzymes that have incomplete descriptions or several distinct functions, motivating the need for more powerful annotation techniques that can predict functionality from sequence alone.

We first visualized the global structure of PAIR representations for EC number prediction. EC number is a hierarchical classification scheme where each digit represents a progressively finer classification of the enzyme function. This allows us to construct a tree-like structure of EC numbers, where sibling nodes sharing the same parent should have related functions. We calculated the normalized Hausdorff distance between the EC number clusters defined by the baseline protein representation (e.g., ProtT5) and our enhanced representation (e.g., PAIR(ProtT5)). The Hausdorff distance measures the separation between two sets of points. Figure 5c displays a heatmap of the pairwise Hausdorff distances normalized by the maximal distance, with darker shades indicating greater dissimilarity between the compared EC clusters. We ordered the EC numbers using a depth-first traversal of the EC tree, grouping together numbers with common parents. The heatmap shows that PAIR exhibits higher contrast between the diagonal and off-diagonal cells. This suggests PAIR has encoded a more well-defined structure that better captures enzyme functional similarity relationships.

**Figure 5:**
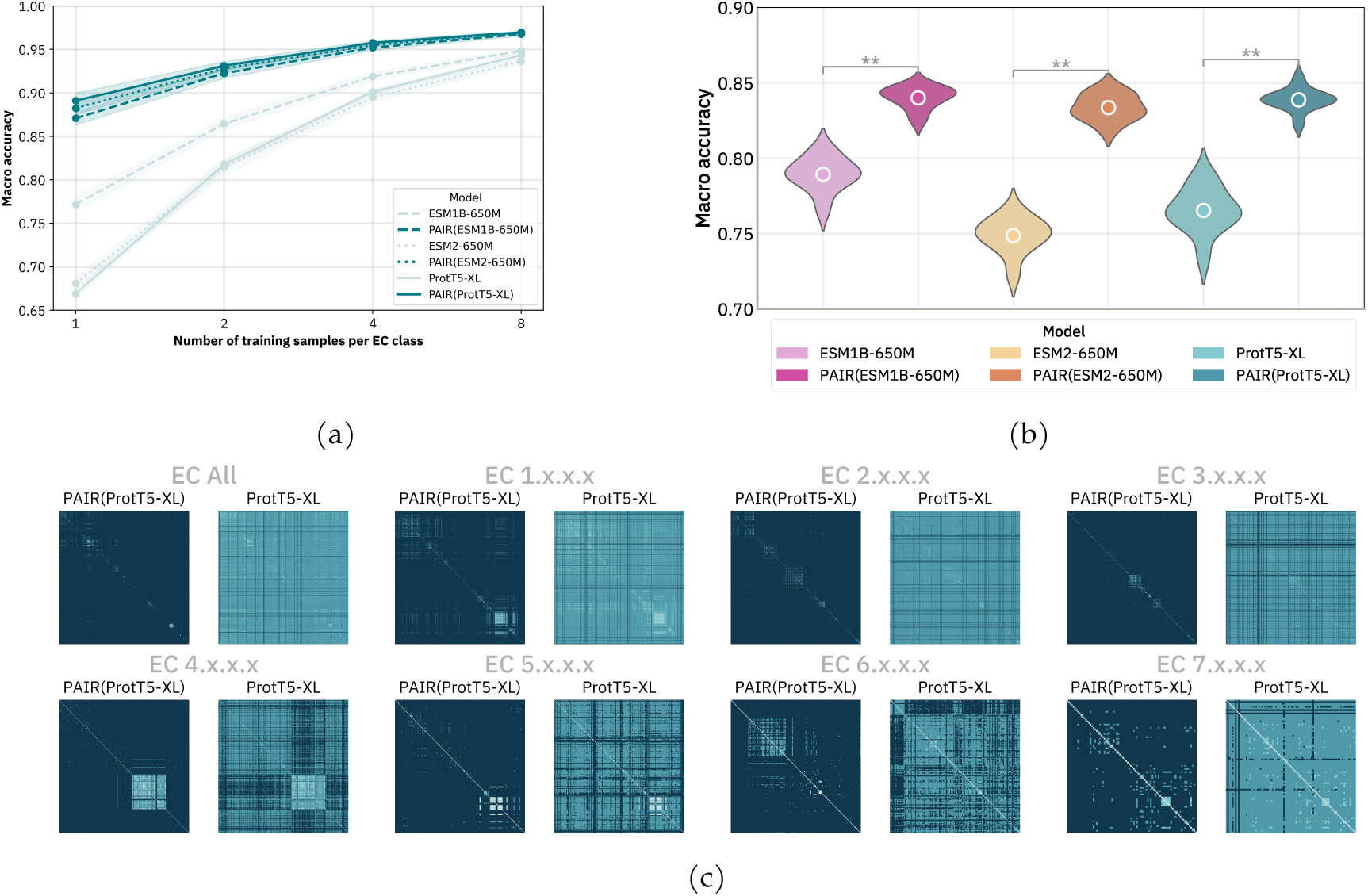
PAIR is better at capturing enzyme function similarity. (a)**PAIR improves few-shot EC classifications.** We train a linear classifier on top of the frozen embeddings with 1, 2, 4, or 8 training samples per EC class. The light blue line represents the base protein models and the dark blue line represents the PAIR-enhanced models. PAIR-enhanced models outperform the baseline models across all shots. The gap between the PAIR and baseline models becomes larger as the number of training examples decreases. (b) **PAIR improves one-shot EC classifications for low-resource enzymes.** We train a linear classifier on 1 training sample per EC class for low-resource EC numbers (those that have 3-9 samples in the entire dataset). We repeat each experiment for 10 random seeds. We find that the PAIR outperforms each baseline protein language model significantly (** indicates *p*-value *<* 0.02 according to a Wilcoxon signed-rank test). (c) **PAIR has a more distinct cluster structure that represents enzyme function relationships.** The EC number classification hierarchically groups enzymes by function, allowing construction of a tree where related nodes indicate similar catalytic activities. We computed the normalized Hausdorff distance between EC clusters defined by ProtT5 and PAIR(ProtT5). The darker shade of the cell in Figure 5c indicates more dissimilar cluster pairs. PAIR (ProtT5) has a higher contrast between the colors of diagonal and off-diagonal cells.

Motivated by this, we further investigated the ability of our model to predict enzyme functionality. We fine-tuned a linear classifier on top of PAIR and baseline model embeddings to predict EC numbers in a few-shot setting. We selected enzymes from our Swiss-Prot 2023 dataset that were annotated with a single EC number and belonged to EC classes with at least 10 members. We randomly split them into train (80%), validation (10%) and test (10%) subsets. For each EC class in the training set, we sampled *k* training samples, where *k ∈ {*1, 2, 4, 8*}* and fine-tuned a linear classifier to predict the enzyme class. We fine-tuned the learning rate on the validation set and plotted the performance on the test set in Figure 5a. We repeated this 10 times for each shot and plotted the standard deviation as error bars. We find that at all *k* training shots, there is a clear separation over PAIR embeddings compared to baseline models. Even with 1 training sample per EC class, PAIR embeddings achieve at least 87.1 *±* 0.8% accuracy, compared to 67-77% for the baseline models. As the number of training shots increases, performance for all models goes up, but PAIR embeddings consistently maintain superior performance.

In practice, many EC classes are understudied, i.e., have very few annotations, and thus predicting them can become more challenging. Therefore, we further investigated the performance of one-shot learning with PAIR embeddings in this (low-resource) scenario. We considered the subset of Swiss-Prot 2023 containing EC classes with 3-9 members. We then split them into training (1 sample per EC class), validation (1 sample per EC class) and test (remaining samples) sets and fine-tuned a linear classifier to predict the EC number. We show the performance on the test set in Figure 5b, comparing PAIR embeddings to embeddings from their corresponding baseline. We find that even in low-resource settings, PAIR embeddings obtain close to 85% accuracy on average, which is significantly higher than baseline models (*p*-value < 0.02). This demonstrates that functional annotations can significantly enhance the performance of protein embeddings, especially in scenarios where training data is limited.

## 3 Discussion

We introduced PAIR: a flexible fine-tuning framework to improve the quality of protein representations by leveraging a diverse set of text annotations. These annotations directly contain information about the fundamental functional properties of proteins. Combining them resulted in a substantial improvement on the protein representation’s ability to predict a variety, including unseen, functional properties of new proteins.

Notably, PAIR consistently outperformed the base protein language model when annotating crucial functional properties for previously uncharacterized proteins. We also found that the embedding space produced by PAIR better captured enzymatic functions, enabling accurate predictions of enzyme commission numbers from just a few examples. In comparison to BLAST, PAIR either matched or exceeded BLAST’s performance across all tasks while being more computationally efficient. Furthermore, PAIR demonstrates even larger improvements over BLAST when the predictions require extrapolation: the tested sequences have low sequence similarity with the training data.

Our work suggests that protein models can be improved by incorporating additional modalities beyond sequence data. One promising direction is to integrate 3D structural information, genomic data and function annotations together into pretraining. Integrating more data modalities could produce even richer and more generalizable protein representations. Furthermore, the flexible nature of PAIR makes it amenable to extensions beyond just proteins. It would be interesting to adapt this framework to learn representations for other biological entities like small molecules, nucleic acids, or even higher-order molecular complexes. Effectively representing and reasoning about the diverse components within biological systems could open up new frontiers for more general-purpose biological models.

## 4 Methods

### 4.1 Model

#### 4.1.1 Model architecture

We used the attention-based sequence-to-sequence architecture proposed in Vaswani et al. [50]. The encoder and the decoder are initialized by the pretrained models. A randomly-initialized cross-attention is then added between the encoder and the decoder, which enables the decoder to gather relevant information from each position of the input sequence. Our implementation follows `transformers.EncoderDecoderModel` from Huggingface [54].

The core building block of the sequence-to-sequence model is the attention mechanism. The attention function maps a query vector *Q* and set of key-value vector pairs (*K, V*) to an output vector. The output is computed as a weighted sum of the value vectors *V*. Each weight is calculated by a similarity function between *Q* and *K*. Vaswani et al. [50] used the scaled dot product as the similarity function:

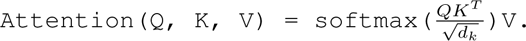

It has proven advantageous to perform multiple attention operations in parallel, called multi-head attention mechanism. Specifically, the queries *Q*, keys *K*, and values *K* are linearly projected *h* times using distinct learned linear transformations. The outputs of these *h* parallel attention computations are then combined, typically through concatenation and linear projection, to yield the final attended representation:

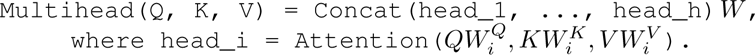

Let *x* = (*x*_1_*, · · ·, x_n_*) denote the input protein sequence and *y* = (*y*_1_*, · · ·, y_m_*) denote the output text annotation. Each *x_i_* is an amino acid and each *y_i_* is a text token. The encoder first maps (*x*_1_*, · · ·, x_n_*) to a sequence of continuous representations *V_N_ ∈* **R***^n×d^* through *N* self-attention blocks, where *n* is the sequence length and *d* is the representation dimension. Self-attention is a special attention mechansim where the query, key and value all come from the previous layer of the encoder *V_j−_*_1_. It allows the model to learn the interaction between different amino acids. The self-attention computation at layer *j* is defined as:

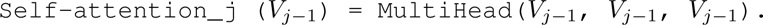

After the encoder processes the protein sequence, the representation of the last layer *V_N_* is then passed to the decoder to generate the text annotation *y* in an auto-regressive fashion. The training objective is to maximize the probability of predicting the next token correctly for each position:

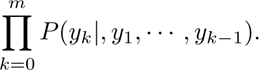

Different from the encoder, the attention block of the decoder consists of a self-attention layer followed by a cross-attention layer. The self-attention layer is similar to the encoder except that it operates on the prefix (*y*_1_*, · · ·, y_k−_*_1_). It means the decoder cannot attend to the tokens after the current position *k*. After the self-attention layer, the representations *U_k_* are then passed to the cross-attention to incorporate the information from the encoder. In cross-attention, the queries of the attention come from the previous self-attention layer *U_k_*, and the keys and values come from the output of the encoder *V_N_*:

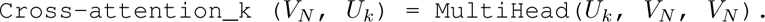

Note that in the case where the encoder and the decoder output representation of different lengths, we add a linear projection head on the encoder side to map its representation to the same dimension as the decoder.

After going through a stack of attention blocks, the decoder then outputs a continuous representation *r_k_* for the position *k*. We then projected *r_k_* using a linear head to compute the logits over the vocabulary of the decoder. At the end, we minimized the cross-entropy loss between this logit and one-hot encoding of the true token *y_k_*to maximize *P* (*y_k_|, y*_1_*, · · ·, y_k−_*_1_).

#### 4.1.2 Pretrained Models

##### Protein Encoder

We studied pretrained protein models from two model families: ProtT5 [16] and ESM [39] [28]. ProtT5 adopts the architecture of T5 and the training objective of BERT [12]. ProtT5 is trained on all sequences in UniRef50 [47]and BFD100 [45]. We only used the encoder component of ProtT5v with 1.5 billion parameters.

We picked two model variants of ESM: ESM4541b [39] and ESM2-650M [28]. ESM2-650M and ESM4541b use similar model architectures and training methodologies. They differ in some training details and time stamps of the training set. ESM uses the same architecture as BERT [14] and is trained using the masked language modelling objective on all 250 million sequences in UniParc [11]. ESM has shown great success in various protein tasks, such as predicting the secondary structure [29] and mutation effect [19].

##### Text Decoder

We initialized the decoder with SciBERT [7]. SciBERT used the same architecture as BERT [14] and was trained on a random sample of 1.14 million papers from Semantic Scholar. 82% of the papers in SciBERT’s training corpus are from the biomedical domain, where most of our annotations are from. SciBERT also trained its own tokenizer on the scientific corpus.

#### 4.1.3 Training Details

We trained all models on the 14 annotation types: domain, family, recommended name, alternative name, function, binding sites, keywords, active sites, cofactors, sites, subunit, PTM, paper title and sucellular location. In order to save memory, we trained our model in the half-precision format of bfloat16 [23]. bfloat16 is a low-precision data format specifically designed for deep learning to improve the speed and memory cost of training while maintaining the performance as full-precision training. Our training is performed on an HPC cluster of AMD M100 GPUs. We used `torch.nn.parallel.DistributedDataParallel` to enable multi-node training. We trained 2 ESM models on 4 nodes, and ProtT5 on 8 nodes. Each node consist of eight 32GB M100 GPUs.

During the training process, we followed a two-step sampling approach for each annotation. First, we sampled the type of annotation, with a higher probability given to annotation types that covered a more diverse range of unique UniRef50 protein clusters. Secondly, within the sampled annotation type, we sampled specific protein annotations with a probability inversely proportional to the size of their UniRef50 cluster. In other words, proteins from smaller UniRef50 clusters (presumably with more unique structures) were preferentially sampled. This downsampling of larger, more redundant clusters helped reduce bias toward highly similar protein structures in the training data. For the protein sequence, we chopped the sequence from a random position if it’s longer than 1024. For the text annotation, we truncate the token length to 128.

We tuned hyperparameters for each model based on the performance of the nine evaluation tasks on the validation set. Afterwards, we retrained the models with the best hyperparameters on the whole February-2023 Swiss-Prot dataset. For the final model, we used the Adam optimizer [32] with learning rate 1*e −* 4 and weight decay 1*e −* 4. We clipped the gradient norm to 1. Our total batch size across all GPUs is 160 for ESM and 128 for ProtT5. We trained ESM2-650M and ESM4541b for 50000 steps, and ProtT5 for 30000 steps.

### 4.2 Evaluation

#### 4.2.1 Function annotations for uncharacterized proteins

Below, we present a brief description of each task, how we parsed the label(s), and what we used as evaluation metrics.

##### Gene Ontology annotations (GO)

GO is a structured knowledgebase developed to unify the representation of genes and gene product attributes (including proteins) across all species. It provides a controlled, hierarchical vocabulary to describe three aspects of protein annotations: molecular function (GO-MF), biological process (GO-BP), and cellular component (GO-CC). We parsed GO labels from the `<feature type=“GO”>` tag in the downloaded Swiss-Prot XML. Similar to Kulmanov and Hoehndorf [25], we only kept the annotations with tags denoting that the labels were validated from real-world experiments. Our evaluation uses a weighted *F*_1_ score following Kulmanov and Hoehndorf [25].

##### Enzyme Commission number (EC)

The EC numbering system is a classification mechanism for the enzymes based on the chemical reactions they catalyze. It consists of 4 digits, each of which represents a progressively more refined categorization. We obtained the EC numbers from the tags of `<feature type=“EC”>`. Then we removed any numbers with fewer than 4 digits. We also removed numbers if they contained the character “n”, which indicates that the annotation has not yet been officially assigned. We used *F*_1_ as the evaluation metric for this task.

##### Recommended protein name (Name)

Gane et al. [17] showed that if the protein name can be predicted, it can give information about its function. Inspired by this, we extracted the recommended name of the protein from the `<protein>/<recommendedName>/<fullName>` tag. For evaluation, we computed the exact match accuracy.

##### UniProt protein family (Family)

A protein family is a set of proteins with common evolutionary origin. UniProt assigns a family name based on information from external protein family databases, sequence similarity and analysis tools, and literature. This is documented in the “Sequence similarities” section on UniProt, and typically appears as a sentence *“Belongs to the XX family”*. We parsed the family name from the `<comment type=“similarity”>/<text>` tags and computed the exact match accuracy during evaluation.

##### Pfam domains (Domain)

A protein domain is a distinct structural and functional unit within the protein. We extracted the Pfam domain annotations from `<dbReference type=“Pfam”>/<property type=“entry name”>` tags. We used *F*_1_ as the evaluatiion metric.

##### Active sites

Active sites are related to the enzymatic behaviour and describe which protein components are involved in catalytic reactions. On UniProt, this information exists as a table where one column is a residue number and another describes the residue role (e.g. “proton acceptor”, “eletrophile”). We extracted the residue roles from the `<feature type=“active site”>` tags in the XML file. We used *F*_1_ as the evaluation metric.

##### Binding sites

This task studies the interaction between proteins and small ligands, such as metals and cofactors. We extracted the names of the ligands within `<feature type=“binding site”> <ligand>` tags and used *F*_1_ as the evaluation metric.

#### 4.2.2 External benchmarks

##### Binary localization [3]

This is a binary classification task that predicts whether a protein is “membrane-bound” or “soluble”. We followed the training (5161), test (1746), validation (1727) split by Xu et al. [55]. The dataset is split to test the model’s generalization behaviour across homologous proteins. The protein sequences undergo a clustering process based on a 30% sequence identity threshold, after which they are partitioned into five distinct subsets. Four of these subsets are utilized for training and validation, while the remaining fold is reserved for test. The evaluation metric is accuracy.

##### Subcellular Localization [3]

This is a more fine-grained version of Binary localization. The task asks to classify the locations of a protein in a cell into ten categories: *cell membrane*, *cytoplasm*, *endoplasmic reticulum*, *golgi apparatus*, *lysosome*, *mitochondrion*, *nucleus*, *peroxisome*, *plastid*, *extracellular*. We used the same the split of Xu et al. [55] (training: 8945, val: 2248, test: 2768). The split was conducted in a similar way as binary localization to test the model’s ability to correctly predict the subcellular localization of homologous proteins. The evaluation metric is accuracy.

##### Fold [21]

The task aims to aims to categorize the global structural topology of a protein into one of the 1194 distinct fold designations. We used the original split in Hou et al. [21] (training: 12312, validation: 736, test: 718). To construct the test set, entire superfamilies are held out from the training and validation. This approach enables an assessment of the model’s capability to identify proteins exhibiting similar structural topology despite dissimilar sequences, i.e. remote homology detection. The evaluation metric is accuracy.

##### Drug Target Interaction [31] [13]

The effect of a small-molecule drug is primarily determined by its binding affinity with the target protein. The drug-target interaction prediction task seeks to computationally predict the interaction activity score between a protein sequence and a small molecule. We used two datasets for this task: DAVIS [13] and BindingDB [31]. Both datasets are regression tasks that predict the equilibrium dissociation constant (*K_D_*) of a protein-molecule pair. The smaller the *K_D_* value, the greater the binding affinity of that pair is. We downloaded both datasets from Therapeutics Data Commons [22]. To evaluate our models, we used the Pearson correlation coefficient with the *K_D_*value following Huang et al. [22].

We applied cold-start split [22] on the protein sequences to split the datasets into training/validation/test with a ratio of 6 : 2 : 2. The cold split is a split method for multi-instance prediction problems that involve two types of entities such as protein and molecules. We first partitioned data based on only the protein sequences into training, validation, and test sets. Then, all pairs associated with the entities in each set are correspondingly assigned to the respective training, validation, and test splits. This split makes sure that there are no overlapped protein sequences across different partitions, so we can evaluate the model’s generalization for unseen proteins.

#### 4.2.3 Baselines

##### Naive

The simplest baseline is to assign the annotation with the highest frequency in the training set to every test protein. For GO, we followed [58] and assigned each protein the top 100 GO terms.

##### BLAST

Basic Local Alignment Search Tool (BLAST) is a widely-used sequence alignment tool that allows users to compare an input or query sequence against a predefined database of sequences to find the most similar one [4]. BLAST uses heuristics on local protein regions to approximate the Smith-Waterman algorithm in order to increase the speed of alignment. For each query, BLAST returns similar sequences in the database above a specified significance threshold (E-value), which represents the number of hits with a similar alignment score that could be found by chance in a random database of a similar size. We used BLASTp, which is specialized for comparing protein sequences. We downloaded the BLASTp executable ^1^. For each downstream task, we created a BLAST database from sequences in Swiss-Prot 2023-02 that had a label for the given task: `makeblastdb -in swissprot202302_{task}.fasta -out {task}_database -dbtype prot`. For each query sequence in the test set, we ran the BLAST algorithm to obtain a list of sequence hits from the database: `blastp-query swissprot202401_{task}.fasta -db {task}_database -evalue 1000`. Note that we set a high E-value to ensure that each query sequence had a hit. We then took the hit with the lowest E-value (most significant) as the closest training sequence and used it for evaluation.

##### ProtST

ProtST is a multi-modal protein language model that was trained to align protein sequences with four types of natural language descriptions from Swiss-Prot (Function, Subcellular Location, Family, Name) [56]. This model was the first to show that pre-training on both sequences and text descriptions can boost performance on downstream protein tasks. We built on this work by incorporating several more annotation categories and reducing the loss function to a simple next token prediction loss, and show that even with a simpler framework, incorporating more annotations results in better performance. We obtained ProtST embeddings from their model checkpoints ^2^

##### ProtBert

ProtBert is a BERT-based protein language model trained on 217 million unlabelled amino acid sequences from UniRef100 using a masked language modelling loss [15]. We obtained the model from HuggingFace ^3^ and followed the same sequence pre-processing. We set the maximum length to 1026 (1024 for amino acids and 2 for special tokens).

##### Galactica

Galactica is a decoder-only Transformer model trained on 48 million scientific papers, textbooks, websites, and other sources of scientific information [48], outperforming several large language models on a range of science and math tasks. We obtained embeddings from Galactica-6.7B ^4^ by using the same prompt format as in Taylor et al. [48]. We set the maximum length to 1026 (1024 for amino acids and 2 for special tokens) and averaged the embeddings from the last hidden state.

##### GPT

GPT models have previously demonstrated domain knowledge in biology and chemistry [1, 8, 35]. To create the most similar settings as ours, we followed [1] and used API calls to obtain embeddings from GPT-3 (`text-embedding-3-large`, the most capable GPT model providing access to embeddings at the time of writing). We structured the prompt using the FASTA format.

## 5 Acknowledgements

We would like to thank the following people for insightful discussions: Cait Harrigan, Andrew Jung and Yangjun Ruan. This work was supported in part by Advanced Micro Devices, Inc. under the AMD AI&HPC Fund program, as well as by the Acceleration Consortium and the Vector Institute.

## A Related Work

### A.1 Computational techniques for protein labelling

Detecting protein sequence homology - common evolutionary ancestry - using sequence similarity is the standard approach to identifying functions that are common between proteins. A common way to do so is determining pairwise alignment across large sets of labeled sequences to retrieve similar protein sequences to a query using the Basic Local Alignment Search Tool (BLAST) algorithm [4]. BLAST is a carefully-designed heuristics-based algorithm that approximates the optimal alignment between sequences. Other methods include Hidden Markov Models (HMMs)[43] and integrate other information like protein family, amino acid composition and evolutionary information [5, 9, 46, 52]. These methods are the state-of-the-art of classical representation methods. While they are reliable for proteins that have high sequence similarity (*>* 25%), they struggle at remote homology detection – accurate classification of sequences that have low similarity to the current databases. This has motivated the emergence of deep learning efforts [26, 21, 3, 27, 42].

### A.2 Protein Language Models

Recently, numerous studies on protein representations learned in an unsupervised manner on large-scale protein sequence corpuses were reported in the literature [40, 15, 2, 38]. Such works usually use existing language model architectures and training objectives from natural language processing (NLP), and are therefore frequently called protein language models (PLMs) [49, 51]. The hope is that biochemical and physicochemical principles that underlie sequence–structure–function relationships can be captured and be used to predict the functional properties of proteins [59, 37, 38, 49, 55].

### A.3 Protein representation learning with text annotations

In an effort to enrich the pretrained protein representations, several works have augmented the training pipeline with modalities such as natural language descriptions. A pioneering work is DeepText2GO, which uses abstracts from the National Library of Medicine and homology information together to predict GO annotations of protein sequences [57]. Another example is the OntoProtein model, which jointly optimizes the embedding of the GO knowledge graph, composed of text descriptions, and the protein sequence embedding using contrastive learning during pre-training [60]. Both works mentioned are trained using a single source of annotations.

The work most closely related to ours is ProtST [56]. ProtST employs a combination of four training objectives: masked protein modeling, InfoNCE contrastive loss, protein-to-text masked language modeling, and text-to-protein masked language modeling. ProtST is trained on four types of annotations: function, name, subcellular location and family. In comparison to ProtST, our approach offers three distinct advantages: 1) We leveraged the entire Swiss-Prot database and trained on a more extensive set of annotation types. 2) Our training loss formulation is significantly simpler, while obtaining superior results. 3) We conducted a more rigorous evaluation using a temporal data split and held-out proteins.

## B Data split

To split the dataset into the training and validation sert, we obtained the 2023-02 UniRef50 [47] cluster centers with members in Swiss-Prot 2023-02. Then, we ran MMseqs2 [44] on the cluster centers using a similar search as Lin et al. [29] with a sequence similarity threshold of 10%: `–min-seq-id 0.1 –alignment-mode 3 –max-seqs 300 -s 7 -c 0.8 –cov-mode 0`. We randomly selected 10% of the resulting reference sequences and generated the validation set by combining the members of those UniRef50 clusters that were present in Swiss-Prot 2023-02. We repeated the same procedure for the remaining 90% of reference sequences to create the training set.

**Table A:** Validation set performance of ESM2-150M fine-tuned with different types of annotations.

|  | EC | Name | GO-CC | GO-MF | GO-BP | Domain | Family | Binding | Active |
| --- | --- | --- | --- | --- | --- | --- | --- | --- | --- |
| ESM2-150M | 0.292 | 0.213 | 0.387 | 0.317 | 0.232 | 0.515 | 0.436 | 0.573 | 0.630 |
| Recommended Name | 0.535 | 0.279 | 0.385 | 0.355 | 0.243 | 0.704 | 0.581 | 0.754 | 0.774 |
| Domain | 0.560 | 0.282 | 0.372 | 0.366 | 0.244 | 0.726 | **0.615** | 0.738 | 0.827 |
| Family | 0.511 | 0.281 | 0.381 | 0.371 | 0.248 | 0.721 | 0.606 | 0.754 | 0.811 |
| Alternative Name | 0.518 | 0.271 | 0.387 | 0.357 | 0.246 | 0.694 | 0.568 | 0.724 | 0.822 |
| Function | 0.527 | 0.278 | 0.385 | 0.360 | 0.239 | 0.703 | 0.584 | 0.746 | 0.828 |
| Keywords | 0.489 | 0.273 | 0.386 | 0.350 | 0.237 | 0.689 | 0.539 | 0.764 | 0.774 |
| Binding sites | 0.527 | 0.274 | 0.372 | 0.352 | 0.238 | 0.697 | 0.592 | 0.756 | 0.798 |
| Active sites | 0.455 | 0.265 | 0.366 | 0.343 | 0.233 | 0.648 | 0.529 | 0.683 | 0.800 |
| Sites | 0.359 | 0.245 | 0.369 | 0.322 | 0.227 | 0.581 | 0.478 | 0.624 | 0.670 |
| Cofactor | 0.501 | 0.266 | 0.365 | 0.341 | 0.233 | 0.661 | 0.546 | 0.709 | 0.742 |
| Paper title | 0.311 | 0.211 | 0.394 | 0.323 | 0.232 | 0.502 | 0.425 | 0.599 | 0.608 |
| subunit | 0.425 | 0.271 | 0.384 | 0.340 | 0.236 | 0.637 | 0.538 | 0.629 | 0.708 |
| PTM | 0.326 | 0.223 | 0.382 | 0.303 | 0.225 | 0.509 | 0.435 | 0.604 | 0.610 |
| Subcellular location | 0.297 | 0.221 | **0.403** | 0.323 | 0.231 | 0.499 | 0.431 | 0.571 | 0.621 |
| Induction text | 0.229 | 0.182 | 0.370 | 0.268 | 0.214 | 0.423 | 0.362 | 0.508 | 0.542 |
| Biophysical | 0.284 | 0.198 | 0.375 | 0.294 | 0.216 | 0.464 | 0.398 | 0.573 | 0.572 |
| Activity regulation | 0.267 | 0.193 | 0.368 | 0.286 | 0.211 | 0.439 | 0.382 | 0.542 | 0.567 |
| Tissue | 0.231 | 0.181 | 0.368 | 0.276 | 0.208 | 0.419 | 0.357 | 0.507 | 0.529 |
| Organism | 0.102 | 0.105 | 0.361 | 0.252 | 0.202 | 0.262 | 0.228 | 0.357 | 0.376 |
| Combined annotations | **0.568** | **0.296** | 0.383 | **0.375** | **0.249** | **0.737** | 0.606 | **0.766** | **0.818** |

**Table B:** Temporal test-set performance of different models.

| Model | EC | Name | GO-CC | GO-MF | GO-BP | Domain | Family | Binding | Active |
| --- | --- | --- | --- | --- | --- | --- | --- | --- | --- |
| Naive | 0.003 | 0.000 | 0.354 | 0.185 | 0.189 | 0.001 | 0.000 | 0.115 | 0.247 |
| BLAST | **0.577** | **0.147** | 0.396 | **0.425** | 0.304 | 0.855 | 0.665 | 0.652 | 0.745 |
| ESM2-650M | 0.464 | 0.118 | 0.395 | 0.338 | 0.281 | 0.722 | 0.563 | 0.598 | 0.669 |
| ESM4541b | 0.522 | 0.129 | 0.378 | 0.373 | 0.296 | 0.809 | 0.622 | 0.670 | 0.709 |
| ProtT5 | 0.513 | 0.128 | 0.398 | 0.363 | 0.288 | 0.801 | 0.604 | 0.641 | 0.706 |
| Prot-BERT | 0.392 | 0.088 | 0.350 | 0.251 | 0.229 | 0.565 | 0.427 | 0.530 | 0.586 |
| Galactica | 0.134 | 0.047 | 0.254 | 0.169 | 0.162 | 0.203 | 0.174 | 0.235 | 0.406 |
| GPT | 0.068 | 0.030 | 0.222 | 0.126 | 0.110 | 0.099 | 0.099 | 0.162 | 0.307 |
| ProtST(ESM2-650M) | 0.506 | 0.126 | 0.409 | 0.337 | 0.295 | 0.787 | 0.615 | 0.616 | 0.695 |
| ProtST(ESM1b) | 0.547 | 0.131 | 0.384 | 0.364 | 0.297 | 0.841 | 0.658 | 0.679 | 0.726 |
| **PAIR(ESM2-650M)** | 0.554 | 0.134 | **0.442** | 0.374 | 0.290 | 0.847 | 0.648 | 0.679 | 0.752 |
| **PAIR(ESM1b)** | 0.561 | 0.142 | 0.402 | 0.409 | **0.314** | 0.862 | 0.662 | 0.695 | **0.772** |
| **PAIR(ProtT5)** | 0.574 | 0.146 | 0.415 | **0.425** | 0.312 | **0.869** | **0.674** | **0.697** | 0.765 |

**Figure A:**
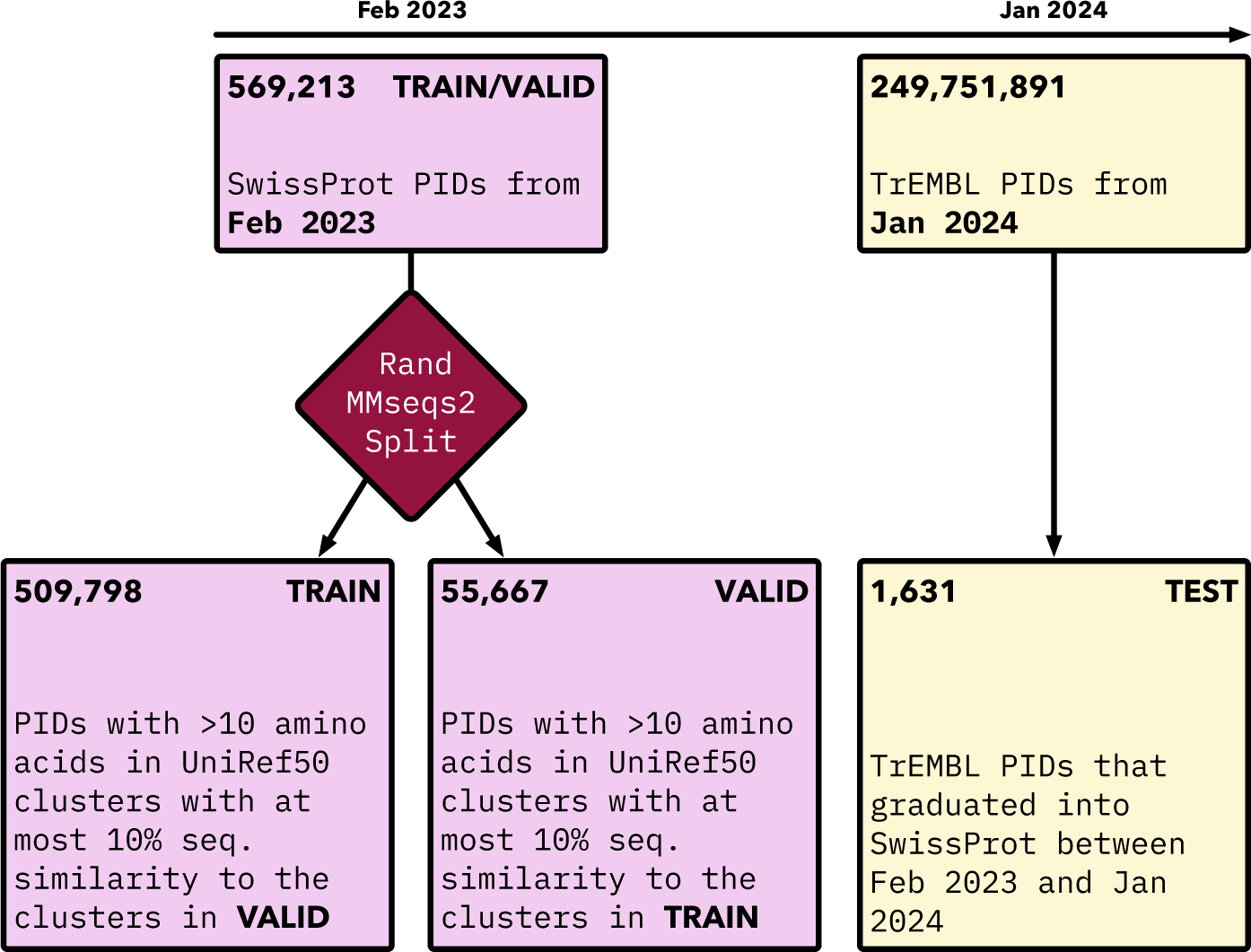
Data splits for training, validation, and test sets. Training and validation sets were created from the February 2023 checkpoint of SwissProt by doing a 10% structural split on UniRef50 cluster centers with MMseqs2. A temporal test set was created by taking the proteins that were added to SwissProt between February 2023 and January 2024.

## C Data Parsing

In the following subsections, we describe how we parsed various annotations of UniProt. We downloaded a snapshot of the Swiss-Prot XML from 2023-03 (train/validation). We parsed the raw XML tree once before training. During training, we generated a data buffer by sampling a set of random proteins and a random annotation category, and then reformatted the annotations in that category to a text prompt. Each data point was added to the buffer as a dictionary containing the following information:

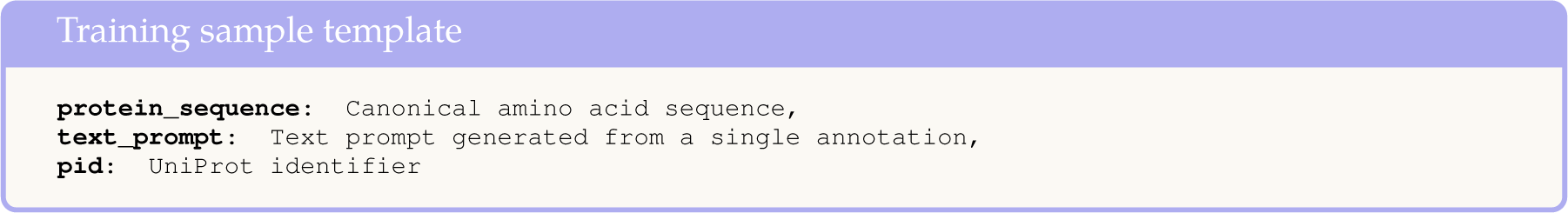

For a given batch, dictionaries from the buffer were randomly sampled. From each dictionary, we passed the value of the “protein_sequence” key to the protein encoder and use the value of the “text_prompt” key as the target output during training. In each paragraph below, we describe how we parsed and loaded each annotation category.

### Function

The function section on UniProt, which describes any information related to the general function of the protein described in natural text. To parse the function section, we extracted the contents of all `<comment type=“function”> <text>` tags in the UniProt XML and added them to a list. During training, we loaded the list of function descriptions. For each description, we first removed any PubMed references. We then remove any sentences that contain the prefix “(By similarity)”, as those are inferred from sequence similarity. Finally, we anonymized the description by replacing any instances of the protein name with “this protein” to prevent the model from falsely achieving high performance simply by memorizing the protein name. We found that for each function description, the first sentence is often a summary of the entire paragraph. Because descriptions could be very long (making it more difficult for the model to learn), we extracted only the first sentence of each function paragraph and added each to the data buffer separately. An example of a training sample containing function information is:

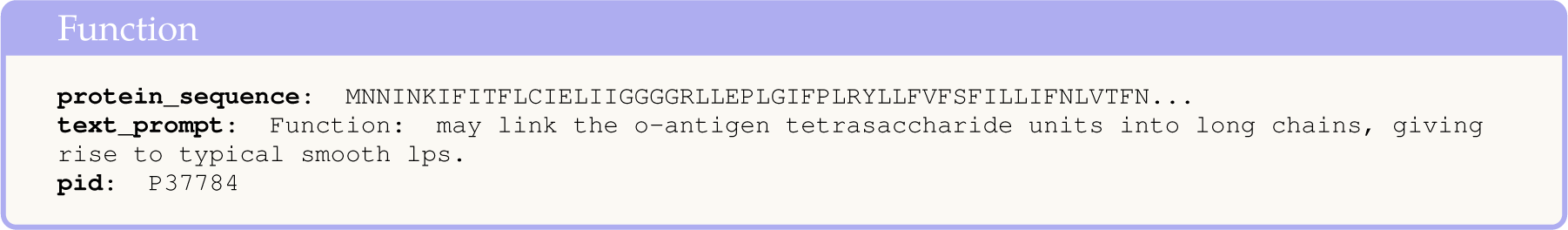

### Active sites

This fact type relates to enzymatic behaviour and describes protein sections that are involved in catalytic reactions. On UniProt, this fact type exists as a table where one column is a residue number and another describes the residue role (e.g. proton acceptor, eletrophile). We extracted the name description of the active sites and their locations from the `<feature type=“active site”>` tags in the XML file and created a dictionary of `{residue_role: [residue_1, residue_2]}`. During training, we appended all the active sites to the data buffer:

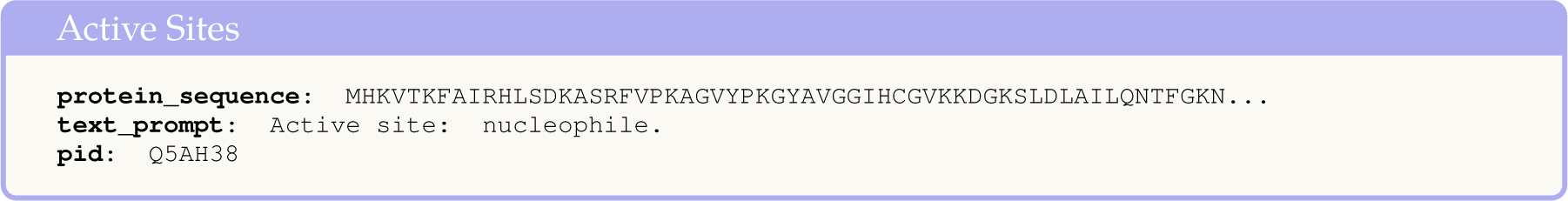

### Binding sites

This fact type describes the interaction between protein residues and small ligands, such as metals, cofactors, and regulators. On UniProt, this fact type exists as a table, with information about the binding site type and location. During parsing, we extracted the names and locations contained within `<feature type=“binding site”> <ligand>` tags in the XML file and generated a dictionary where the key is the ligand and the values are a list of amino acid locations where that ligand binds. During training, we sampled a random protein and looped over all of its binding sites, adding each of them independently to the data buffer using the following format:

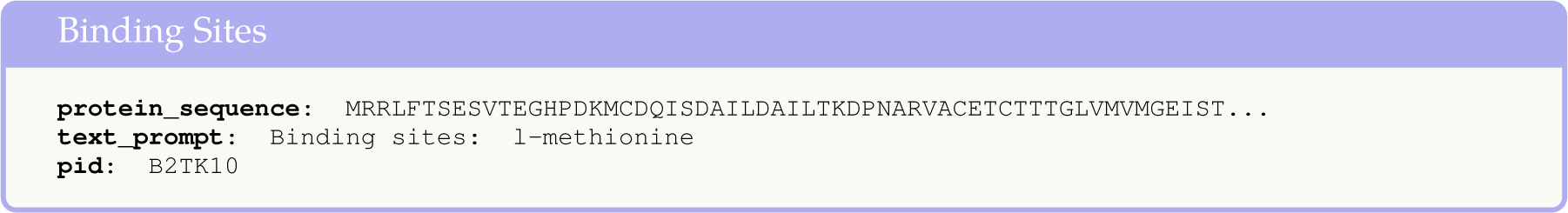

### Sites

The sites section on UniProt refers to any notable single amino acid sites on the protein that are not active sites or binding sites, such as cleavage sites or inhibitory sites. Similar to active sites, we extracted the name description and location information of all `<feature type=“site”>` tags and stored is as as a dictionary, where the keys are the site names and values are locations. During training, we loaded these annotations using the structure of the following example:

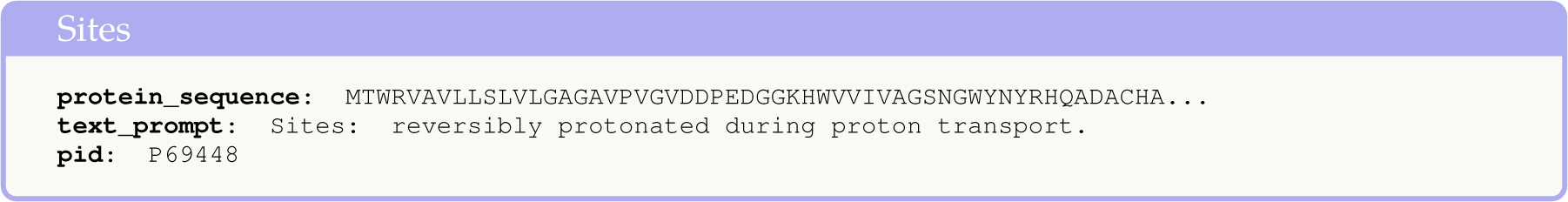

### Activity regulation

The activity regulation section on UniProt describes mechanisms for regulating the activity of enzymes, transporters, and transcription factors, through both activation or inhibition. This section exists as natural language sentences. For a given protein, we extracted the contents of all `<comment type=“activity regulation”> <text>` tags on UniProt. If there were multiple descriptions for a given protein identifier, we combined them into a single description. During training, we first cleaned the description to remove any PubMed identifiers and added it to the buffer:

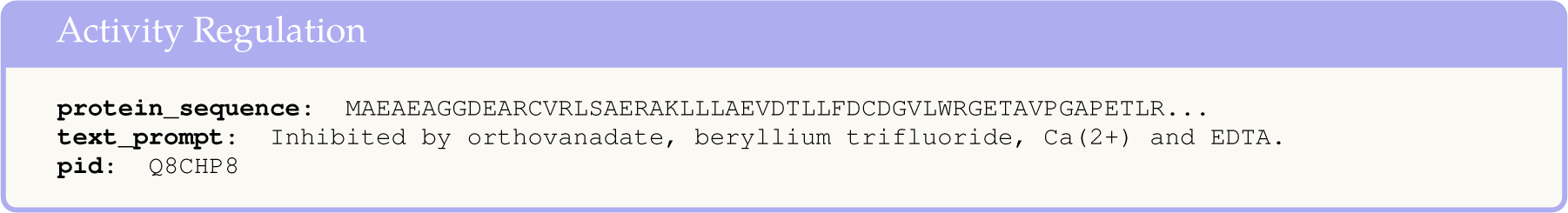

### Biophysicochemical properties

This section describes chemical and physical properties of proteins, including: reaction kinetics, light absorption, redox potentials, and dependence on temperature and pH. We extracted all tags that are children of `<comment type=“biophysicochemical properties”>` from the XML (for example, `<comment type=“biophysicochemical properties”>/<phDependence>, <comment type=“biophysicochemical properties”>/<temperatureDependence>`). We then removed any PubMed identifiers and added them to a list. Most biophysicochemical properties were natural text, but we noticed that absorption and redox potential tended to only be the numerical value and unit, and so we added the prefixes “Absorption: “ and “Redox potential: “ for those properties to provide more context. During training, we added each biophysicochemical property value to the data buffer separately:

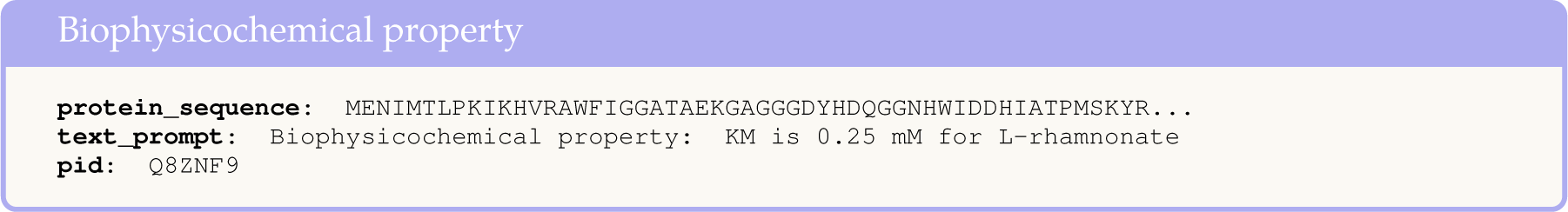

### Cofactors

This section describes any non-protein entities that are required by the protein to engage in catalytic activity, such as metal atoms or vitamins. Cofactors were parsed by extracting the contents of the `<comment type=“cofactor”>/<cofactor>/<name>` tags and adding them all to a list. During traing, each cofactor name was added independently to the data buffer using the following format:

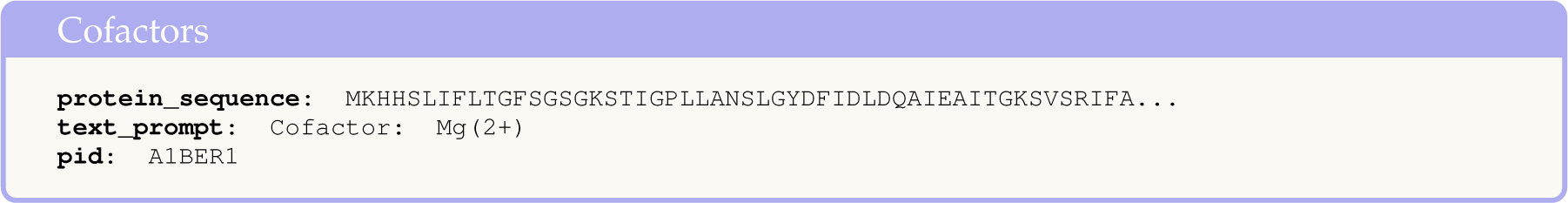

### Domains

In UniProt, the Domains section describes domain(s) present in a protein. The subsection can provide both natural descriptions about the general roles of domains in the protein, as well as a list of domains from various external databases, such as Pfam ^5^. We extracted the contents of all `<comment type=“domain”>/<text>` tags to obtain any natural language descriptions, as well as all values in `<dbReference type=“Pfam”>/<property type=“entry name”>` tags for a given protein and stored them in a list. Because the Pfam domains are abstracted as codes (e.g. “APP_E2”), we added a step when loading the data to map the Pfam code to its natural language description, obtained from UniProt documentation ^6^. For example, the natural language description of “APP_E2” is “E2 domain of amyloid precursor protein”. When loading the data, we looped over each domain in the list (both Pfam and natural description) and added it separately to the data buffer:

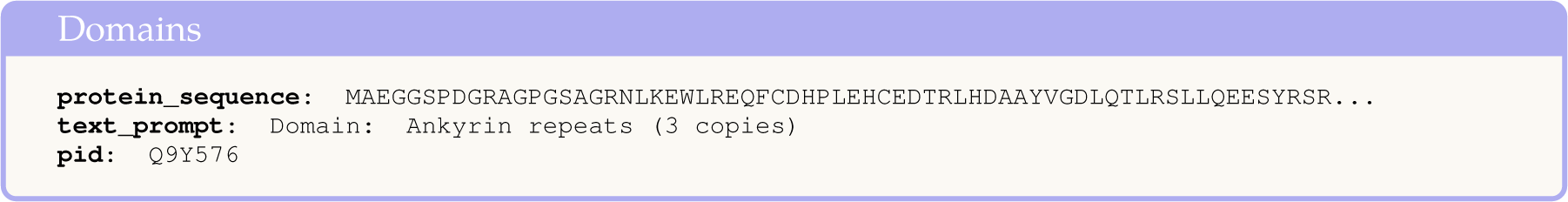

### UniProt protein family

This subsection identifies the protein family (or families) that the protein belongs to. UniProt assigns a family name based on information from external protein family databases (such as InterPro ^7^), sequence similarity and analysis tools, and literature. This is documented in the “Sequence similarities” section on UniProt, and typically appears as a sentence *“Belongs to the XX family”*. We parsed the family name from the `<comment type=“similarity”>/<text>` tags from the UniProt XML. In the event of multiple instances, we joined them into a single string. During training, we removed any PubMed IDs from the string and loaded it into the buffer.

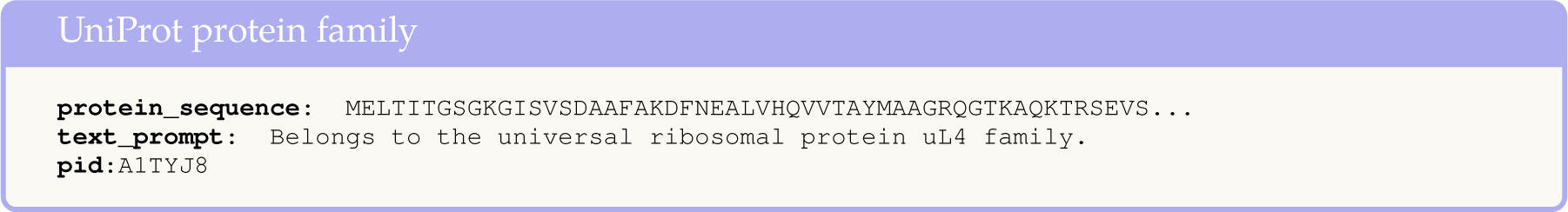

### Keywords

UniProt identifies the most relevant keywords that summarize each protein. These keywords are from a controlled vocabulary, which are structured into ten higher-level categories: *Biological process, cellular component, coding sequence diversity, developmental stage, disease, domain, ligand, molecular function, post-translational modification, technical term*. During parsing, we extracted keywords for each protein from the `<keyword>` tags, along with the keyword ID. We obtained a mapping of each keyword ID to its higher-level category ^8^. When loading the data during training, we only included keywords if they did not belong to the following categories: ‘Technical term’, ‘Developmental stage’, ‘Biological process’, ‘Cellular component’, ‘Molecular function’. We excluded ‘Technical term’ and ‘Developmental stage’ since they are not related to protein function and added noise to the labels. We removed ‘Biological process’, ‘Cellular component’, ‘Molecular function’ since these are related to GO annotations, which are downstream tasks we evaluated on.

We added each keyword to the data buffer individually:

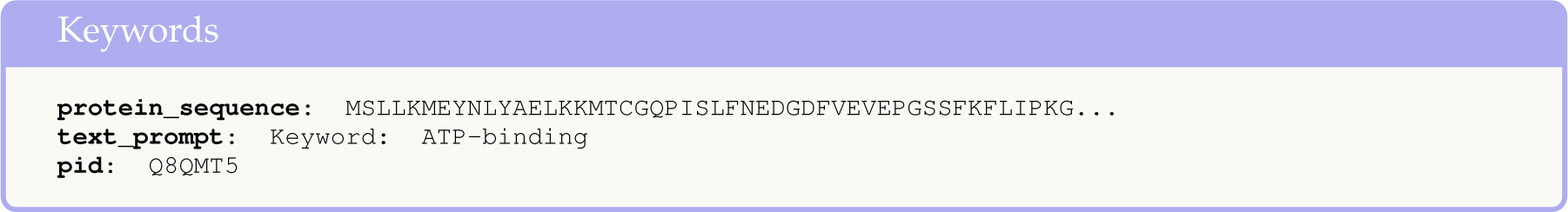

### Tissue specificity

Tissue specificity describes where mRNA and proteins are expressed in cells or tissue (if the organism is multicellular) in natural language. We extracted the contents of all `<comment type=“tissue specificity”>/<text>` tags and joined all descriptions into a single string. When loading this annotation during training, we removed PubMed references from the description and anonymized it from the protein name. Below is an example:

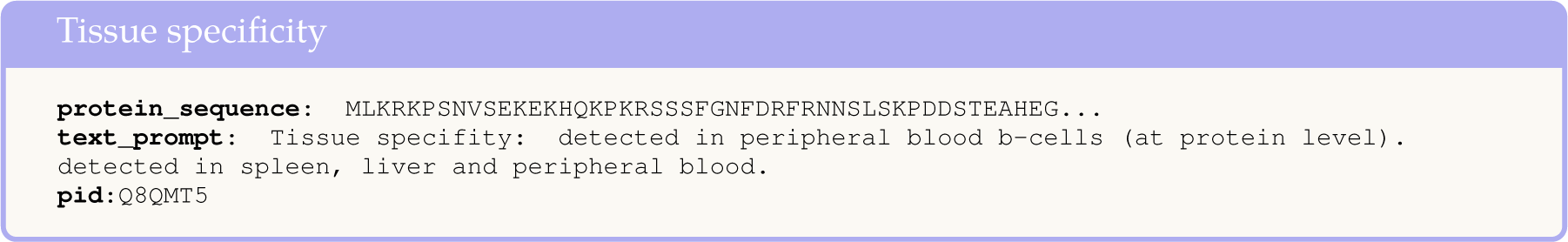

### Organism

Organism is a subsection of the protein taxonomy that identifies the name of the organism the protein was sourced from. The organism name can consists of the Latin scientific name, followed by the common English name, or only the common name in the case of viruses. To parse the organism name, we extracted the values of the `<organism>/<name type=“common”>` and `<organism>/<name type=“scientific”>` tags. When loading the data, we only kept common organism names. If the name contained information about a bacterial or viral strain (e.g. Influenza A virus (strain A/Aichi/2/1968 H3N2)), we removed information related to the strain since we did not expect our model to be able to learn it well.

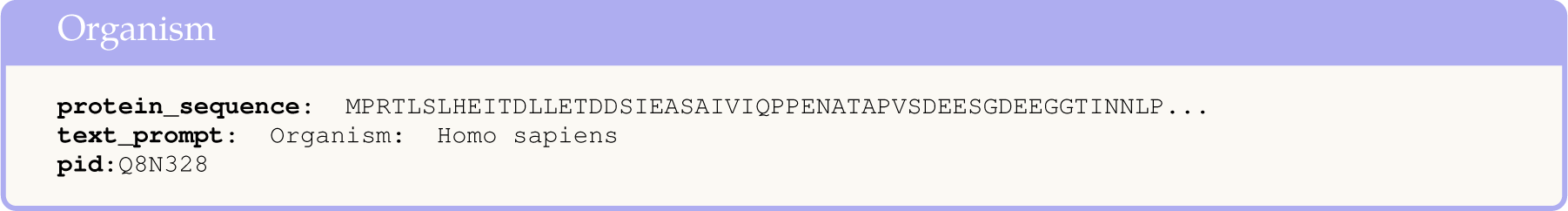

### Paper titles

The motivation of using paper titles as an annotation category is that they can provide a summary of important findings related to a protein. The UniProt XML contains paper titles in `<reference>/<citation>/<title>` tags, as well as an attribute related to the scope of the paper (e.g. FUNCTION, EXTRACELLULAR COPPER-BINDING DOMAIN) in `<reference key=“1”>/<scope>` tags. We extracted all paper titles and scopes for a protein and added them to a list. During training, we added each paper title separately to the buffer if its scope contained the word “FUNCTION”:

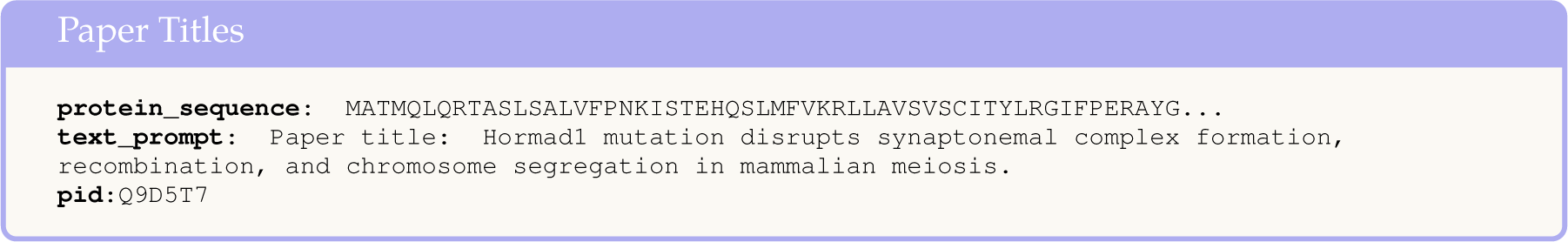

### Recommended names

Gane et al. [17] showed that if the protein name can be predicted, it can give information about its function. Inspired by this, we extracted the Recommended name of the protein, which is the official protein name agreed upon by the UniProt consortium. We extracted the name from the `<protein>/<recommendedName>/<fullName>` tag. During training, we added it to the data buffer as shown below:

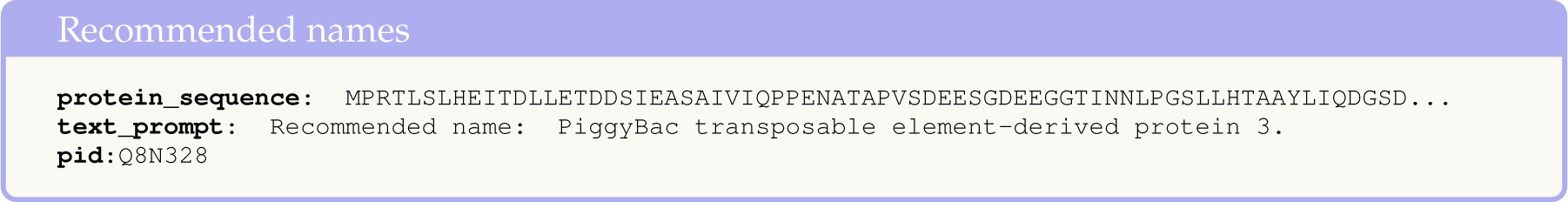

### Alternative names

We also included alternative names of the protein, which are synonyms of the recommended name. We extracted all synonyms from the `<alternativeName>/<fullName>` tags. During training, we loaded the list of alternative names for a given protein and joined them into a single prompt:

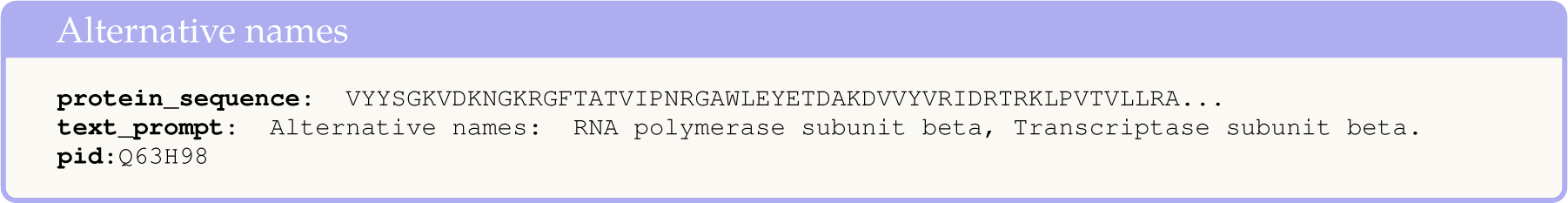

### Subunits

Subunits describe quaternary structures of a protein, as well as how they interact with other proteins (protein-protein interactions) in natural language. We extracted the contents of each `<comment type=“subunit”>/<text>` tag in the UniProt XML and added each description to a list. When loading the data, we removed PubMed references and the “(By similarity)” suffix from each description, and add the first sentence of each description as separate entries into the data buffer. Below is an example:

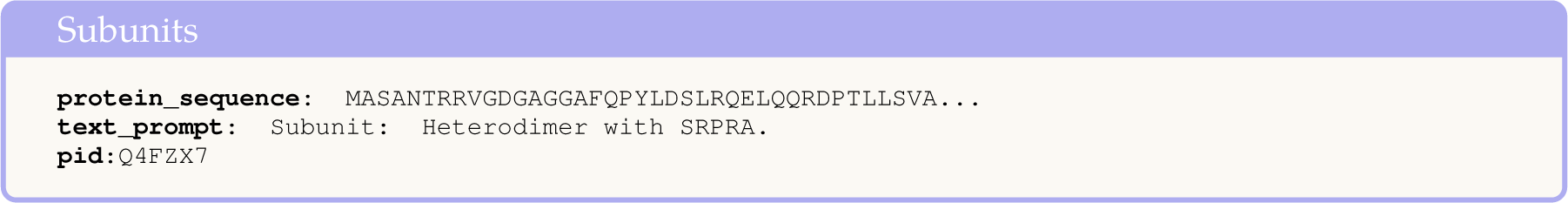

### Post-translational modifications

Post-translational modifications (PTMs) describe covalent modifications to amino acids, which can modulate the function of a protein. We extracted the PTMs that are described using natural language on UniProt by parsing the contents of the `<comment type=“PTM”>/<text>` tags. During loading, we extracted the first sentence of the description, anonymized it, removed PubMed substrings, removed any instances of the phrase “(By similarity)”, and added it to the data buffer.

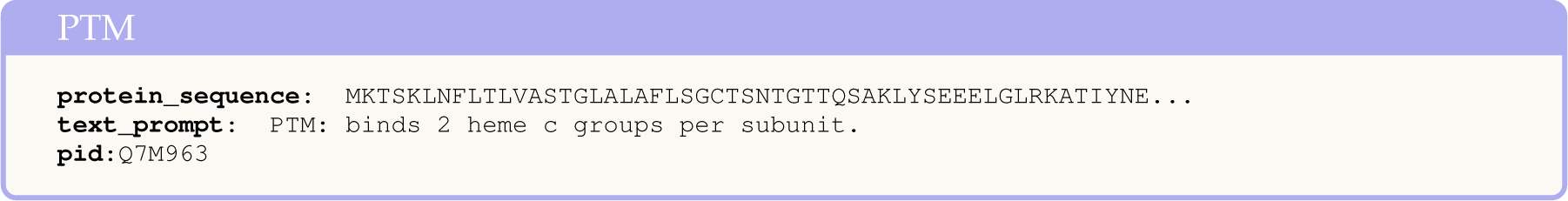

### Subcellular location

This section on UniProt describes the location of the protein in the cell. We extracted the contents of all `<comment type=“subcellular location”>/<subcellularLocation>/<location>` and `<comment type=“subcellular location”>/<subcellularLocation>/<topology>` tags, as well as any natural language descriptions in `<comment type=“subcellular location”>/<text>` and added them to separate lists (one for each tag). For each protein, we generated a dictionary with keys “locations”, “topologies”, and “text”, and set the correspondings lists as values. During data loading, we passed all locations, topologies, and text descriptions separately into the data buffer.

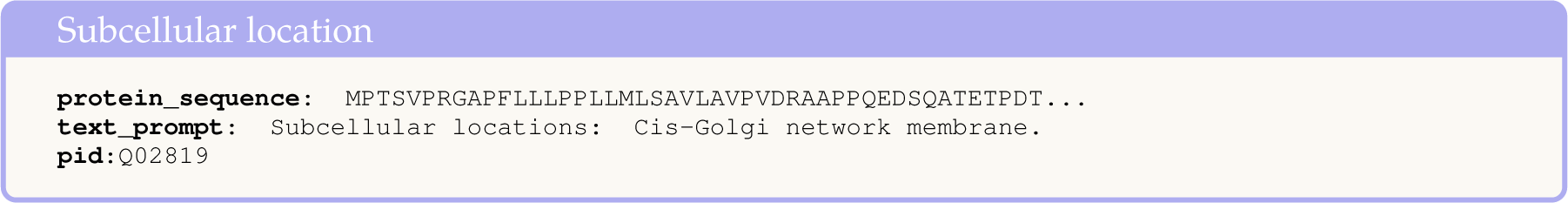

### Induction

The Induction section is a natural language description of how a protein can be upregulated or down-regulated in the presence of inducers or repressors like chemical compounds. We extracted all natural language descriptions from `<comment type=“induction”>/<text>` tags into a single prompt. During data loading, we removed PubMed substrings from the prompt and anonoymized it before adding it to the buffer.

**Figure B:**
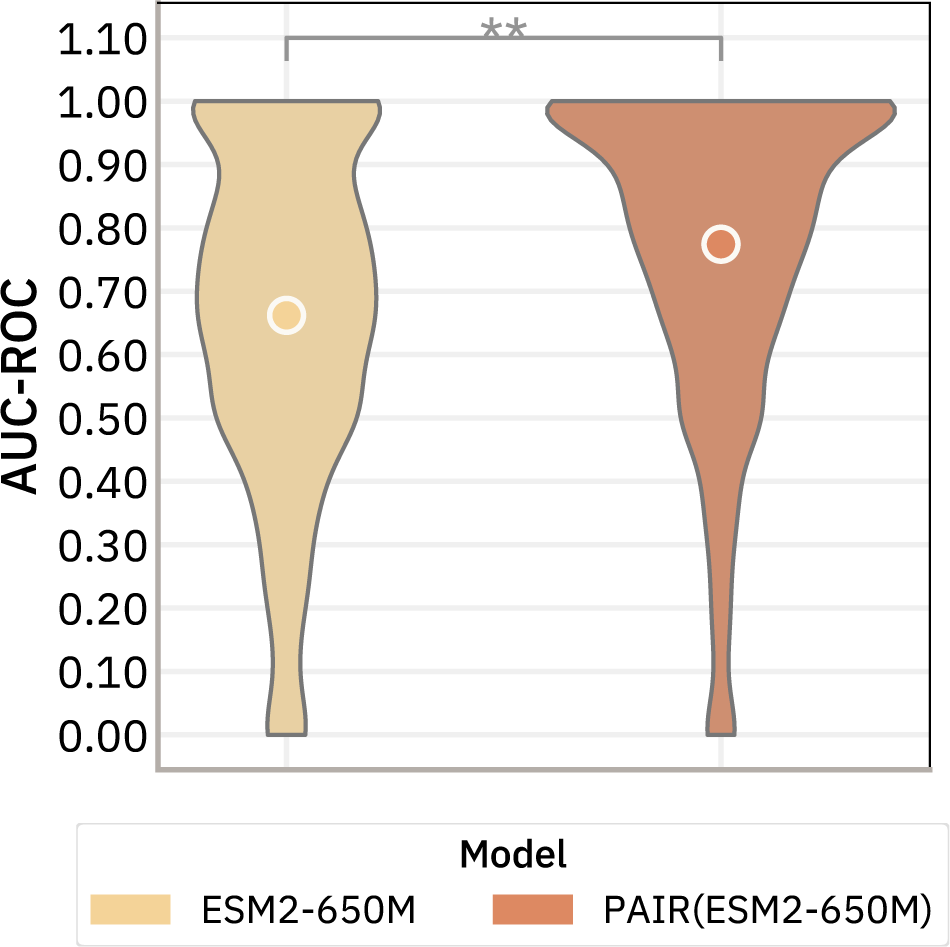
PAIR improves mutation effect predictions. We plot the distribution of the AUC-ROC scores for ESM2-650M and PAIR(ESM2-650M) over assays. The result shows that PAIR improve the mutation effect predictions (** indicates *p*-value *<* 0.02 according to a Wilcoxon signed-rank test).

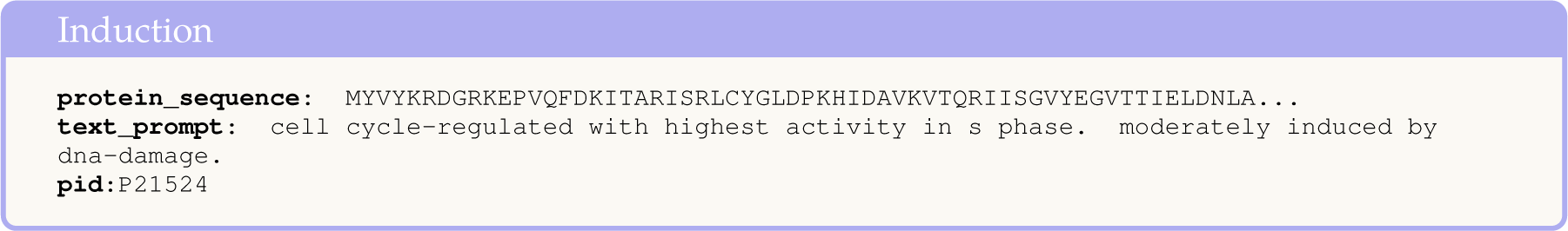

## D Protein Mutation Effects

The protein mutation effect task aims to predict whether amino acid substitutions in proteins are likely to be benign or pathogenic. We focus on the model’s ability to distinguish between these two classes of mutations based on their protein sequence representations. Specifically, we used the *l*_2_ distance between the representations of the canonical sequence and each variant sequence to predict the mutation effect. The underlying hypothesis is that a high-quality representation model should map benign sequences closer to the canonical sequence than pathogenic ones in the learned feature space.

We compared ESM2-650M and PAIR(ESM2-650M) on the clinical substitution benchmark in ProteinGym [36]. The dataset consists of 2050 clinicaly-annotated assays and around 65*k* human protein mutation labels. To quantify the model’s discriminative ability, we calculated the Area Under the Receiver Operating Characteristic curve (AUC-ROC) score of the representation distance. Figure B plots the distribution of the AUC-ROC scores over all assays for ESM2-650M and PAIR(ESM2-650M). We find that PAIR(ESM2-650M) outperforms ESM2-650M in predicting mutation effects. This result is particularly intriguing, considering that our model’s training data did not specifically include variant information or mutation-related tasks. We think this implies our representations more closely resemble the function space.

## Footnotes

1 https://ftp.ncbi.nlm.nih.gov/blast/executables/blast+/LATEST/

2 https://github.com/DeepGraphLearning/ProtST

3 https://huggingface.co/Rostlab/prot_bert

4 https://huggingface.co/facebook/galactica-6.7b

5 http://pfam.xfam.org

6 https://ftp.uniprot.org/pub/databases/uniprot/previous_major_releases/release-2023_02/knowledgebase/,Pfam-A.clans.tsv

7 https://www.ebi.ac.uk/interpro/

8 https://ftp.uniprot.org/pub/databases/uniprot/previous_major_releases/release-2023_02/knowledgebase/,keywlist.txt

## References

[1] Microsoft Research AI4Science and Microsoft Azure Quantum. The impact of large language models on scientific discovery: a preliminary study using gpt-4. arXiv preprint arXiv:2311.07361, 2023.

[2] Ethan C Alley, Grigory Khimulya, Surojit Biswas, Mohammed AlQuraishi, and George M Church. Unified rational protein engineering with sequence-based deep representation learning. Nature methods, 16(12):1315–1322, 2019.

[3] José Juan Almagro Armenteros, Casper Kaae Sønderby, Søren Kaae Sønderby, Henrik Nielsen, and Ole Winther. Deeploc: prediction of protein subcellular localization using deep learning. Bioinformatics, 33(21):3387–3395, 2017.

[4] Stephen F Altschul, Warren Gish, Webb Miller, Eugene W Myers, and David J Lipman. Basic local alignment search tool. Journal of molecular biology, 215(3):403–410, 1990.

[5] Stephen F Altschul, Thomas L Madden, Alejandro A Schäffer, Jinghui Zhang, Zheng Zhang, Webb Miller, and David J Lipman. Gapped blast and psi-blast: a new generation of protein database search programs. Nucleic acids research, 25(17):3389–3402, 1997.

[6] Iz Beltagy, Kyle Lo, and Arman Cohan. Scibert: A pretrained language model for scientific text. arXiv preprint arXiv:1903.10676, 2019.

[7] Iz Beltagy, Kyle Lo, and Arman Cohan. Scibert: A pretrained language model for scientific text. arXiv preprint arXiv:1903.10676, 2019.

[8] Andres M Bran, Sam Cox, Oliver Schilter, Carlo Baldassari, Andrew D White, and Philippe Schwaller. Chemcrow: Augmenting large-language models with chemistry tools. arXiv preprint arXiv:2304.05376, 2023.

[9] Kuo-Chen Chou. Using amphiphilic pseudo amino acid composition to predict enzyme subfamily classes. Bioinformatics, 21(1):10–19, 2005.

[10] Dimitrios Christofidellis, Giorgio Giannone, Jannis Born, Ole Winther, Teodoro Laino, and Matteo Manica. Unifying molecular and textual representations via multi-task language modelling. Proceedings of the 40th International Conference on Machine Learning, 202:6140–6157, 2023. URL https://proceedings.mlr.press/v202/christofidellis23a.html.

[11] The UniProt Consortium. UniProt: the universal protein knowledgebase in 2021. Nucleic Acids Research, 49(D1):D480–D489, 11 2020. ISSN 0305-1048. doi: 10.1093/nar/gkaa1100. URL 10.1093/nar/gkaa1100.

[12] Zihang Dai, Zhilin Yang, Yiming Yang, Jaime Carbonell, Quoc V Le, and Ruslan Salakhutdinov. Transformer-xl: Attentive language models beyond a fixed-length context. arXiv preprint arXiv:1901.02860, 2019.

[13] Mindy I Davis, Jeremy P Hunt, Sanna Herrgard, Pietro Ciceri, Lisa M Wodicka, Gabriel Pallares, Michael Hocker, Daniel K Treiber, and Patrick P Zarrinkar. Comprehensive analysis of kinase inhibitor selectivity. Nature biotechnology, 29(11):1046–1051, 2011.

[14] Jacob Devlin, Ming-Wei Chang, Kenton Lee, and Kristina Toutanova. Bert: Pre-training of deep bidirectional transformers for language understanding. arXiv preprint arXiv:1810.04805, 2018.

[15] Ahmed Elnaggar, Michael Heinzinger, Christian Dallago, Ghalia Rehawi, Yu Wang, Llion Jones, Tom Gibbs, Tamas Feher, Christoph Angerer, Martin Steinegger, et al. Prottrans: Toward understanding the language of life through self-supervised learning. IEEE transactions on pattern analysis and machine intelligence, 44(10):7112–7127, 2021.

[16] Ahmed Elnaggar, Michael Heinzinger, Christian Dallago, Ghalia Rehawi, Yu Wang, Llion Jones, Tom Gibbs, Tamas Feher, Christoph Angerer, Martin Steinegger, et al. Prottrans: Toward understanding the language of life through self-supervised learning. IEEE transactions on pattern analysis and machine intelligence, 44(10):7112–7127, 2021.

[17] Andrea Gane, Maxwell L. Bileschi, David Dohan, Elena Speretta, Amélie Héliou, Laetitia Meng-Papaxanthos, Hermann Zellner, Eugene Brevdo, Ankur Parikh, Maria J. Martin, Sandra Orchard, UniProt Collaborators, and Lucy Colwellm. Protnlm: Model-based natural language protein annotation, 2023. URL https://www.uniprot.org/help/ProtNLM.

[18] Tymor Hamamsy, James T Morton, Robert Blackwell, Daniel Berenberg, Nicholas Carriero, Vladimir Gligorijevic, Charlie EM Strauss, Julia Koehler Leman, Kyunghyun Cho, and Richard Bonneau. Protein remote homology detection and structural alignment using deep learning. Nature biotechnology, pages 1–11, 2023.

[19] Brian L Hie, Varun R Shanker, Duo Xu, Theodora UJ Bruun, Payton A Weidenbacher, Shaogeng Tang, Wesley Wu, John E Pak, and Peter S Kim. Efficient evolution of human antibodies from general protein language models. Nature Biotechnology, 42(2):275–283, 2024.

[20] Thomas A Hopf, John B Ingraham, Frank J Poelwijk, Charlotta PI Schärfe, Michael Springer, Chris Sander, and Debora S Marks. Mutation effects predicted from sequence co-variation. Nature biotechnology, 35(2):128–135, 2017.

[21] Jie Hou, Badri Adhikari, and Jianlin Cheng. Deepsf: deep convolutional neural network for mapping protein sequences to folds. Bioinformatics, 34(8):1295–1303, 2018.

[22] Kexin Huang, Tianfan Fu, Wenhao Gao, Yue Zhao, Yusuf Roohani, Jure Leskovec, Connor W Coley, Cao Xiao, Jimeng Sun, and Marinka Zitnik. Therapeutics data commons: Machine learning datasets and tasks for drug discovery and development. Proceedings of Neural Information Processing Systems, NeurIPS Datasets and Benchmarks, 2021.

[23] Dhiraj Kalamkar, Dheevatsa Mudigere, Naveen Mellempudi, Dipankar Das, Kunal Banerjee, Sasikanth Avancha, Dharma Teja Vooturi, Nataraj Jammalamadaka, Jianyu Huang, Hector Yuen, et al. A study of bfloat16 for deep learning training. arXiv preprint arXiv:1905.12322, 2019.

[24] Siddharth Karamcheti, Suraj Nair, Ashwin Balakrishna, Percy Liang, Thomas Kollar, and Dorsa Sadigh. Prismatic vlms: Investigating the design space of visually-conditioned language models. In International Conference on Machine Learning (ICML), 2024.

[25] Maxat Kulmanov and Robert Hoehndorf. Deepgozero: improving protein function prediction from sequence and zero-shot learning based on ontology axioms. Bioinformatics, 38 (Supplement_1):i238–i245, 2022.

[26] Maxat Kulmanov, Mohammed Asif Khan, and Robert Hoehndorf. Deepgo: predicting protein functions from sequence and interactions using a deep ontology-aware classifier. Bioinformatics, 34(4):660–668, 2018.

[27] Yu Li, Sheng Wang, Ramzan Umarov, Bingqing Xie, Ming Fan, Lihua Li, and Xin Gao. Deepre: sequence-based enzyme ec number prediction by deep learning. Bioinformatics, 34(5):760–769, 2018.

[28] Zeming Lin, Halil Akin, Roshan Rao, Brian Hie, Zhongkai Zhu, Wenting Lu, Nikita Smetanin, Robert Verkuil, Ori Kabeli, Yaniv Shmueli, et al. Evolutionary-scale prediction of atomic-level protein structure with a language model. Science, 379(6637):1123–1130, 2023.

[29] Zeming Lin, Halil Akin, Roshan Rao, Brian Hie, Zhongkai Zhu, Wenting Lu, Nikita Smetanin, Robert Verkuil, Ori Kabeli, Yaniv Shmueli, et al. Evolutionary-scale prediction of atomic-level protein structure with a language model. Science, 379(6637):1123–1130, 2023.

[30] Shengchao Liu, Yutao Zhu, Jiarui Lu, Zhao Xu, Weili Nie, Anthony Gitter, Chaowei Xiao, Jian Tang, Hongyu Guo, and Anima Anandkumar. A text-guided protein design framework. arXiv preprint arXiv:2302.04611, 2023.

[31] Tiqing Liu, Yuhmei Lin, Xin Wen, Robert N Jorissen, and Michael K Gilson. Bindingdb: a web-accessible database of experimentally determined protein–ligand binding affinities. Nucleic acids research, 35(suppl_1):D198–D201, 2007.

[32] Ilya Loshchilov and Frank Hutter. Decoupled weight decay regularization. In International Conference on Learning Representations, 2018.

[33] Minh-Thang Luong, Quoc V Le, Ilya Sutskever, Oriol Vinyals, and Lukasz Kaiser. Multi-task sequence to sequence learning. arXiv preprint arXiv:1511.06114, 2015.

[34] Wayne P. Maddison and Richard G. FitzJohn. The Unsolved Challenge to Phylogenetic Correlation Tests for Categorical Characters. Systematic Biology, 64(1):127–136, 09 2014. ISSN 1063-5157. doi: 10.1093/sysbio/syu070. URL 10.1093/sysbio/syu070.

[35] Adrian Mirza, Nawaf Alampara, Sreekanth Kunchapu, Benedict Emoekabu, Aswanth Krishnan, Mara Wilhelmi, Macjonathan Okereke, Juliane Eberhardt, Amir Mohammad Elahi, Maximilian Greiner, et al. Are large language models superhuman chemists? arXiv preprint arXiv:2404.01475, 2024.

[36] Pascal Notin, Aaron Kollasch, Daniel Ritter, Lood van Niekerk, Steffanie Paul, Han Spinner, Nathan Rollins, Ada Shaw, Rose Orenbuch, Ruben Weitzman, Jonathan Frazer, Mafalda Dias, Dinko Franceschi, Yarin Gal, and Debora Marks. Proteingym: Largescale benchmarks for protein fitness prediction and design. In A. Oh, T. Neumann, A. Globerson, K. Saenko, M. Hardt, and S. Levine, editors, Advances in Neural Information Processing Systems, volume 36, pages 64331–64379. Curran Associates, Inc., 2023. URL https://proceedings.neurips.cc/paper_files/paper/2023/file/cac723e5ff29f65e3fcbb0739ae91bee-Paper-Datasets_and_Benchmarks.pdf.

[37] Predrag Radivojac, Wyatt T Clark, Tal Ronnen Oron, Alexandra M Schnoes, Tobias Wittkop, Artem Sokolov, Kiley Graim, Christopher Funk, Karin Verspoor, Asa Ben-Hur, et al. A largescale evaluation of computational protein function prediction. Nature methods, 10(3):221–227, 2013.

[38] Roshan Rao, Nicholas Bhattacharya, Neil Thomas, Yan Duan, Peter Chen, John Canny, Pieter Abbeel, and Yun Song. Evaluating protein transfer learning with tape. Advances in neural information processing systems, 32, 2019.

[39] Alexander Rives, Joshua Meier, Tom Sercu, Siddharth Goyal, Zeming Lin, Jason Liu, Demi Guo, Myle Ott, C Lawrence Zitnick, Jerry Ma, et al. Biological structure and function emerge from scaling unsupervised learning to 250 million protein sequences. Proceedings of the National Academy of Sciences, 118(15):e2016239118, 2021.

[40] Alexander Rives, Joshua Meier, Tom Sercu, Siddharth Goyal, Zeming Lin, Jason Liu, Demi Guo, Myle Ott, C Lawrence Zitnick, Jerry Ma, et al. Biological structure and function emerge from scaling unsupervised learning to 250 million protein sequences. Proceedings of the National Academy of Sciences, 118(15):e2016239118, 2021.

[41] Sascha Rothe, Shashi Narayan, and Aliaksei Severyn. Leveraging pre-trained checkpoints for sequence generation tasks. Transactions of the Association for Computational Linguistics, 8:264–280, 2020.

[42] Jae Yong Ryu, Hyun Uk Kim, and Sang Yup Lee. Deep learning enables high-quality and high-throughput prediction of enzyme commission numbers. Proceedings of the National Academy of Sciences, 116(28):13996–14001, 2019.

[43] Johannes Söding. Protein homology detection by hmm–hmm comparison. Bioinformatics, 21 (7):951–960, 2005.

[44] Martin Steinegger and Johannes Söding. Mmseqs2 enables sensitive protein sequence searching for the analysis of massive data sets. Nature biotechnology, 35(11):1026–1028, 2017.

[45] Martin Steinegger and Johannes Söding. Clustering huge protein sequence sets in linear time. Nature communications, 9(1):2542, 2018.

[46] Martin Steinegger, Markus Meier, Milot Mirdita, Harald Vöhringer, Stephan J Haunsberger, and Johannes Söding. Hh-suite3 for fast remote homology detection and deep protein annotation. BMC bioinformatics, 20(1):1–15, 2019.

[47] Baris E Suzek, Yuqi Wang, Hongzhan Huang, Peter B McGarvey, Cathy H Wu, and UniProt Consortium. Uniref clusters: a comprehensive and scalable alternative for improving sequence similarity searches. Bioinformatics, 31(6):926–932, 2015.

[48] Ross Taylor, Marcin Kardas, Guillem Cucurull, Thomas Scialom, Anthony Hartshorn, Elvis Saravia, Andrew Poulton, Viktor Kerkez, and Robert Stojnic. Galactica: A large language model for science. arXiv preprint arXiv:2211.09085, 2022.

[49] Serbulent Unsal, Heval Atas, Muammer Albayrak, Kemal Turhan, Aybar C Acar, and Tunca Doğan. Learning functional properties of proteins with language models. Nature Machine Intelligence, 4(3):227–245, 2022.

[50] Ashish Vaswani, Noam Shazeer, Niki Parmar, Jakob Uszkoreit, Llion Jones, Aidan N Gomez, Lukasz Kaiser, and Illia Polosukhin. Attention is all you need. Advances in neural information processing systems, 30, 2017.

[51] Mai Ha Vu, Rahmad Akbar, Philippe A Robert, Bartlomiej Swiatczak, Geir Kjetil Sandve, Victor Greiff, and Dag Trygve Truslew Haug. Linguistically inspired roadmap for building biologically reliable protein language models. Nature Machine Intelligence, 5(5):485–496, 2023.

[52] Jiawei Wang, Bingjiao Yang, Jerico Revote, Andre Leier, Tatiana T Marquez-Lago, Geoffrey Webb, Jiangning Song, Kuo-Chen Chou, and Trevor Lithgow. Possum: a bioinformatics toolkit for generating numerical sequence feature descriptors based on pssm profiles. Bioinformatics, 33(17):2756–2758, 2017.

[53] Jason Wei, Yi Tay, Rishi Bommasani, Colin Raffel, Barret Zoph, Sebastian Borgeaud, Dani Yogatama, Maarten Bosma, Denny Zhou, Donald Metzler, Ed H. Chi, Tatsunori Hashimoto, Oriol Vinyals, Percy Liang, Jeff Dean, and William Fedus. Emergent abilities of large language models. Transactions on Machine Learning Research, 2022. ISSN 2835-8856. URL https://openreview.net/forum?id=yzkSU5zdwD. Survey Certification.

[54] Thomas Wolf, Lysandre Debut, Victor Sanh, Julien Chaumond, Clement Delangue, Anthony Moi, Pierric Cistac, Tim Rault, Rémi Louf, Morgan Funtowicz, et al. Huggingface’s transformers: State-of-the-art natural language processing. arXiv preprint arXiv:1910.03771, 2019.

[55] Minghao Xu, Zuobai Zhang, Jiarui Lu, Zhaocheng Zhu, Yangtian Zhang, Ma Chang, Runcheng Liu, and Jian Tang. Peer: a comprehensive and multi-task benchmark for protein sequence understanding. Advances in Neural Information Processing Systems, 35:35156–35173, 2022.

[56] Minghao Xu, Xinyu Yuan, Santiago Miret, and Jian Tang. Protst: Multi-modality learning of protein sequences and biomedical texts. arXiv preprint arXiv:2301.12040, 2023.

[57] Ronghui You, Xiaodi Huang, and Shanfeng Zhu. Deeptext2go: improving large-scale protein function prediction with deep semantic text representation. Methods, 145:82–90, 2018.

[58] Ronghui You, Zihan Zhang, Yi Xiong, Fengzhu Sun, Hiroshi Mamitsuka, and Shanfeng Zhu. Golabeler: improving sequence-based large-scale protein function prediction by learning to rank. Bioinformatics, 34(14):2465–2473, 2018.

[59] Tianhao Yu, Haiyang Cui, Jianan Canal Li, Yunan Luo, Guangde Jiang, and Huimin Zhao. Enzyme function prediction using contrastive learning. Science, 379(6639):1358–1363, 2023.

[60] Ningyu Zhang, Zhen Bi, Xiaozhuan Liang, Siyuan Cheng, Haosen Hong, Shumin Deng, Jiazhang Lian, Qiang Zhang, and Huajun Chen. Ontoprotein: Protein pretraining with gene ontology embedding. arXiv preprint arXiv:2201.11147, 2022.

